# Molecular basis of foreign DNA recognition by BREX anti-phage immunity system

**DOI:** 10.1101/2024.04.12.589305

**Authors:** Alena Drobiazko, Myfanwy Adams, Mikhail Skutel, Kristina Potekhina, Mikhail Matlashov, Karen L. Maxwell, Tim R. Blower, Konstantin Severinov, Dmitry Ghilarov, Artem Isaev

## Abstract

Anti-phage systems of the BREX (BacteRiophage EXclusion) superfamily rely on epigenetic DNA methylation to discriminate between the host and invading DNA, but their mechanism of protection remains enigmatic. We demonstrate that in Type I BREX systems, both defense and methylation are based on site-specific DNA recognition by the BrxX (PglX) methyltransferase and require the S-adenosyl methionine cofactor. We present a 2.2-Å cryoEM structure of *Escherichia coli* BrxX bound to target dsDNA, which reveals the molecular details of DNA recognition by BREX and paves the way for rational engineering of BREX specificity. We show that BrxX alone does not support methylation, and BREX activity requires an assembly of a supramolecular BrxBCXZ immune complex. Finally, we present a cryoEM structure of BrxX bound to a phage-encoded inhibitor Ocr that sequesters an inactive dimeric form of BrxX. Together, these results allow us to propose a model of BREX-mediated DNA sensing and anti-phage defense.

## INTRODUCTION

Phages are the most abundant biological entities on Earth and their interaction with bacterial hosts shapes the structure and evolution of microbial communities^1,2^ To withstand phage infection, bacteria developed a broad arsenal of immunity systems, the abundance and diversity of which has been appreciated only in recent years^3–6^. Bacterial anti-phage defense strategies can be broadly classified into three main groups: avoidance of host cells recognition by phages, which often involves limiting the availability of surface receptors^7^; targeting of phage genomes or essential phage processes by innate or adaptive immunity systems of the host^4^, and abortive responses that inhibit the metabolism or even cause death of the infected cell, thus halting phage propagation in the population^8,9^. The function of host innate immunity systems depends on the ability to discriminate between self (host) and non-self (virus) molecules. A range of DNA-targeting systems rely on epigenetic DNA modifications, such as DNA methylation, as a marker of self^10^; a classic example being restriction-modification (R-M) systems present in ∼83% of sequenced prokaryotic genomes^11^. A multitude of recently described immunity systems, such as Dnd^12^, Ssp^13^, Dpd^14^, DISARM^15^, MADS^16^, and BREX^17,18^ also encode DNA modification proteins. The mechanisms of phage defense by these systems are not fully determined and likely are more sophisticated than direct nucleolytic cleavage of non-modified invading DNA^19,20^.

BREX, which is found in ∼7% of prokaryotic genomes, often clusters together with other immunity-related genes in defense islands^11,17,21^. Based on protein composition, BREX systems are classified into 6 types^17^, with Type II BREX previously described and partially characterised as a Pgl (phage growth limitation) system of *Streptomyces*^22,23^. The most widespread Type I BREX employs an R-M like principle: it modifies a cell’s own DNA at specific sites, and the acquisition of BREX-specific modification protects the phage from exclusion^18^. BREX methylation is strand-specific, i.e., only one-strand of the asymmetric recognition site (**GGTA<u>A</u>G** in *E. coli* HS) is methylated. Similar methylation patterns are employed by Type ISP, Type IIL, and Type III R-M systems^24–26^. The asymmetric methylation requires the activity of defense systems to be tightly regulated to avoid cleavage of non-methylated sites emerging after replication of host DNA. For example, Type ISP and III R-M systems require simultaneous recognition of two non-methylated sites on opposite DNA strands, followed by translocation of one (Type III) or both (Type ISP) restriction complexes along the DNA and their subsequent collision to licence DNA cleavage^27,28^. Type IIL restriction enzymes assemble into a multimeric complex whilst remaining bound to their cognate site, and the probability of restriction is determined by the “local density” of non-methylated sites^29^. How BREX systems recognize foreign DNA and avoid self-toxicity associated with the inevitable emergence of non-methylated sites in the replicating host DNA is currently unknown.

With the exception of Type IV, all BREX systems encode BrxX (PglX) N6-adenine-methyltransferases^17,18,30–34^, which are thought to be responsible for DNA modification^18^. Other core BREX proteins include BrxZ (PglZ) belonging to alkaline phosphatase superfamily and homologous to the phosphodiesterase PorX^35^, and BrxC (PglY), an AAA+ ATPase with distant homology to ORC/Cdc6 (Origin Recognition Complex) proteins^36,37^. In addition, Type I BREX systems encode a DNA-binding AAA+ ATPase BrxL^38^, and two small proteins of unknown function BrxA^39^ and BrxB (**Figure 1A**). Type I BREX gene clusters are often associated with genes coding for additional accessory proteins, for example, the WYL-domain transcriptional regulator BrxR^33,40^, which is also found in other defense systems such as CBASS^41^. In addition, a hotspot of diverse immunity genes has been identified inside BREX clusters^42^, suggesting a possible synergy between BREX and other systems active against BREX-resistant phages. For example, Type IV R-M nuclease BrxU encoded by some Type I BREX clusters can restrict phages with modified DNA, which are insensitive to BREX^18,31^.

**Figure 1:**
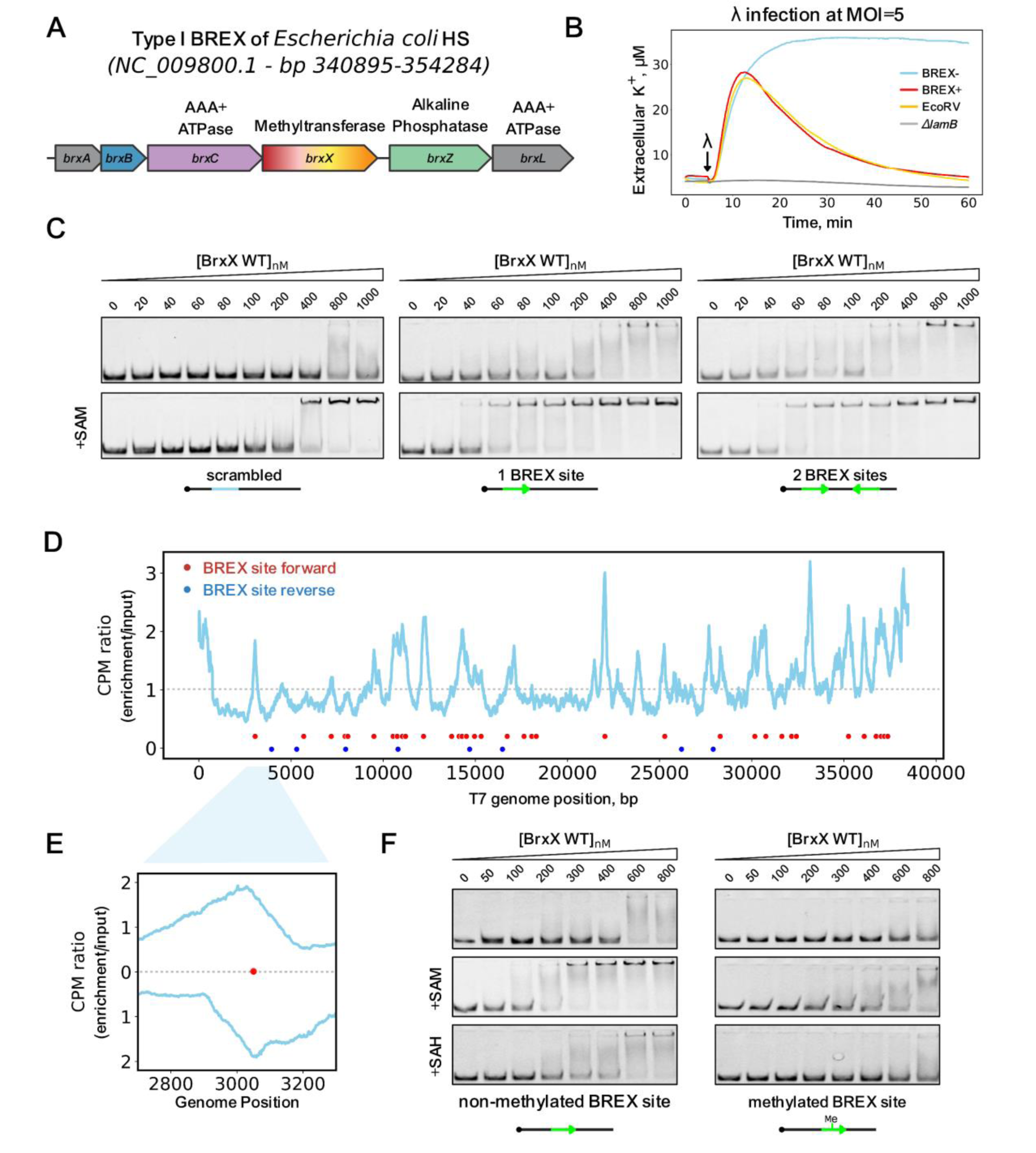
The BrxX (PglX) methyltransferase specifically recognizes BREX sites in DNA. (**A**) The Type I BREX locus of *E. coli* HS, its genomic positions and predicted functions of encoded proteins are shown. (**B**) Potassium efflux assay with *E. coli* BW25113 BREX^-^, BREX^+^, EcoRV, or Δ*lamB* cultures infected with phage *λ* at MOI=5. A representative result from the three independent experiments is shown. (**C**) EMSA with 20 nM Cy5-labelled 43 bp dsDNA substrates with no BREX sites (“scrambled”), or with one or two BREX sites, incubated with an indicated amount of BrxX in the presence or in the absence of 0.5 mM SAM. Green arrows indicate the orientation of BREX sites. Representative gels from experiments performed in triplicate are shown. (**D**) Strep-seq analysis of DNA cross-linked to BrxX-Strep 15 minutes after infection of a BREX^+^ culture with T7fusion phage at MOI=1. Reads from enriched sample were mapped to T7 genome and the signal was normalized to the bulk DNA level and genome size to obtain CPM values. Positions of BREX sites (**GGTA<u>A</u>G**) on the top and bottom DNA strands are indicated with red and blue dots, correspondingly. (**E**) Cross-correlation analysis (normalized coverage of the forward and reverse reads) for a representative peak in the T7 genome. (**F**) EMSA with 20 nM of 40 bp Cy5-labelled dsDNA substrate with methylated or non-methylated BREX site incubated with indicated amounts of BrxX. Representative gels from triplicate experiments are shown.

A model explaining BREX-mediated exclusion of phage DNA is currently lacking. BREX likely acts at the early stages of the infection as it prevents accumulation of newly synthesized phage DNA inside infected cells^17,18^. While BREX methylates host DNA similarly to the R-M systems, it lacks an obvious restriction component. Yet, BREX is inhibited by viral anti-defense proteins that inhibit R-M systems, most notably, a DNA mimic Ocr that interacts with BrxX^43,44^. BREX methylation is also inhibited by the depletion of the intracellular SAM pool by phage T3 S-adenosyl methionine (SAM) lyase. Strikingly, the SAM depletion also inhibits BREX defense, implying that like in some Type I R-M systems, SAM could perform regulatory functions in addition to being a donor of methyl groups^45,46^.

In this work, we directly demonstrate that the BREX response is based on foreign DNA recognition by methyltransferase BrxX. We show that BrxX defines BREX site-specificity, and that DNA binding is enhanced by SAM. We present a 2.2 Å cryoEM structure of BrxX-DNA complex providing a high-resolution structural view of the DNA recognition by this family of enzymes, which highlights that both target recognition and methyltransferase domains are involved in sequence-specific recognition of BREX sites. Using single-particle cryoEM we further demonstrate how DNA mimic anti-restriction protein Ocr of bacteriophage T7 prevents DNA binding by BrxX and locks it in an inactive dimeric state. Finally, we establish that BREX defense and methylation require assembly of a multi-subunit BrxBCXZ complex, allowing us to propose a plausible model of BREX-mediated exclusion of phage DNA to guide further research of this enigmatic system.

## RESULTS

### The BREX defense is based on site-specific recognition of foreign DNA by BrxX

We previously determined that Type I BREX of *E. coli* strain HS methylates the fifth adenine in asymmetric **GGTA<u>A</u>G** motifs in host DNA, and that λ phages that acquire BREX-specific methylation are insensitive to BREX^18^. This implies that the BREX defense requires the recognition of non-methylated BREX sites in infecting phage genomes. We next sought to determine at which stage is BREX defense initiated. Given that the injection of λ DNA into host cells triggers ion efflux^47^, we carried out potassium efflux assay to monitor the dynamics of λ genome injection^48^. We measured extracellular K^+^ concentration in BREX^-^ and BREX^+^ cultures, as well as in cultures of cells harbouring Type II R-M EcoRV infected with λ at high MOI (**Figure 1B**). A *ΔlamB* strain lacking the λ receptor was used as a negative control. The initial efflux dynamic was identical for all cultures, however, the BREX^+^ and EcoRV^+^ cells eventually restored the K^+^ potential, confirming that infection had failed. The results suggest that BREX does not interfere with phage genome injection. Since BREX does not affect phage adsorption^18^, the BREX defense must be initiated by the recognition of non-methylated BREX sites once they appear in the cytoplasm.

To identify component(s) of the BREX system that are responsible for site-specific recognition of non-methylated DNA, we purified each of the six *E. coli* HS BREX proteins from cells transformed with appropriate expression plasmids, and carried out electrophoretic mobility gel shift assays (EMSA) with a 43 bp dsDNA probe carrying a single BREX site. A probe with a scrambled BREX site was used as a control (**Figure 1C, Supplementary Figure S1A**). Consistent with previous observations, BrxA efficiently bound both probes^39^. BrxL also bound both specific and non-specific DNA although with affinity that was lower than previously reported^38^, possibly suggesting that the 43 bp substrate used is too short for efficient binding. Importantly, only BrxX showed enhanced binding to the BREX site-containing probe. This suggests that BrxX is responsible for (at least) two core BREX functions: sensing of non-methylated phage DNA and methylation of host DNA. Consistent with a direct role of BrxX in defense, deletion of *brxX* does not result in toxicity^18^, as would be expected if the remaining components of the BREX system were to attack non-methylated sites in host DNA. To further prove the role of BrxX as a sensor of non-methylated BREX sites, we sequenced DNA cross-linked to BrxX in BREX^+^ culture infected with T7 phage (‘Strep-Seq’ procedure, see ***Methods***). Mapping of reads on phage DNA confirmed the presence of peaks centred around BREX sites on both DNA strands (**Figure 1D**) and demonstrated the expected strand cross-correlation despite the relatively low amplitude of the signal (**Figure 1E**). The result thus confirms that BrxX is involved in sensing of non-methylated BREX sites in phage DNA *in vivo*, the first step in the activation of BREX defense.

### SAM and methylation state modulate the affinity of BrxX to BREX sites

S-adenosyl methionine (SAM) serves as a donor of methyl groups for methyltransferases. SAM also can stabilize DNA-bound protein complexes^45,49^, and was recently shown to be required for the BREX defense^46^. We studied how the presence of SAM affects DNA binding by BrxX (**Figure 1C**). BrxX non-specifically bound the 43-bp dsDNA probe with low affinity. The binding was improved ∼ 3-fold in the presence of SAM (K_d_ ^SAM+^ ∼300 nM compared to K_d_ ^SAM-^ >1 μM). The enhancing effect of SAM on DNA binding was ∼5-fold when the substrate contained a BREX site (K_d_ ^SAM+^ ∼60 nM compared to K_d_ ^SAM-^ ∼300 nM), suggesting that SAM promotes BREX sites recognition/results in more stable BrxX:DNA complexes. To further study BrxX substrate preferences we performed EMSAs with BREX site-containing dsDNA fragments of different lengths. Decreasing dsDNA length resulted in weaker binding, while the presence of a super shifted band at higher BrxX concentrations suggested non-specific binding of additional BrxX molecules to same DNA fragments **(Supplementary Figure S1B**). BrxX also efficiently bound ssDNA substrates, although without a clear preference for the presence of BREX sites, suggesting that bases from both the top and bottom DNA strands are required for recognition (**Supplementary Figure S1C**). To further quantify BrxX interaction with dsDNA, we determined the K_d_ values by an orthogonal approach using biolayer interferometry (BLI) with a biotinylated 40 bp dsDNA probe carrying a single BREX site or the polyC sequence (**Supplementary Figure S2A).** Although BLI confirmed BrxX preference for the DNA containing BREX site in the presence of SAM, an accurate K_d_ estimation was not possible due to the heterogenous binding, likely resulting from interaction of more than one BrxX protein with the dsDNA substrate (**Supplementary Figure S2B,D)**. To overcome this, we measured the K_d_ using a shorter 20 bp dsDNA substrate, which showed weaker binding, however, fit to the 1:1 interaction model. The K_d_ values obtained confirm the stimulating effect of SAM and the preference for BREX site containing DNA, in agreement with the EMSA results (**Supplementary Figure S2C,E**).

Reduced affinity to methylated substrates is a common mechanism of methylation activity control in R-M systems^50,51^. At the post-methylation stage, the methylated adenine is accommodated in the enzyme catalytic pocket together with the S-adenosyl homocysteine (SAH), another product of the methylation reaction. When a SAM-bound methyltransferase binds to methylated DNA, SAM interferes with the correct placement of the methylated adenine in the catalytic pocket. To determine if methylation affects the binding of BrxX to its site, we prepared a pair of identical dsDNA substrates with either methylated or non-methylated BREX site and compared their binding to BrxX in three conditions: without co-factors, in the presence of SAM (to mimic BrxX interaction with methylated host DNA), or in the presence of SAH (to mimic the post-methylation BrxX:DNA complex) (**Figure 1F**). As expected, BrxX had had a much lower affinity for the methylated substrate under all conditions tested. This suggests that: *i)* host DNA methylation prevents recognition by BrxX; *ii)* post-methylation complex is not stable, promoting the release of BrxX and, likely, subsequent exchange of SAH for SAM; and *iii)* methylation sensing by BrxX does not strictly require the presence of a co-factor.

### Overall architecture of the BrxX-DNA complex

We sought to understand the molecular mechanism for site-specific recognition and subsequent methylation of DNA by BrxX. We initially tried X-ray crystallography but failed to obtain any DNA-bound crystals. Cryogenic electron microscopy (cryoEM) analysis of BrxX in the presence of 14 bp and 20 bp DNA oligonucleotides containing a single BREX site was also unsuccessful, with most 2D class averages corresponding to what seemed to be apo BrxX (data not shown). However, when we incubated BrxX with a 43 bp dsDNA fragment containing two BREX sites in a “Head-to-Tail” (H2T) arrangement (DNA H2T; **Figure 2A**) in the presence of SAM, a DNA-bound complex represented the majority of 2D class averages (**Supplementary Figure S3B, Figure 2C**). Single-particle analysis of the 11 152 movie dataset demonstrated the formation of a 1:1 complex (**BrxX_1_:DNA_H2T_**) that was highly homogenous despite the presence of two BREX sites (see **Supplementary Figure S3** for the processing pipeline). Only the first binding site was occupied (**Figure 2A**), suggesting that the presence of DNA downstream of a BREX site is essential for BrxX binding. Despite the relatively small protein size (139 kDa), analysis of **BrxX:DNA** resulted in a 2.2 Å global resolution map **(Figure 2 BC)** with local resolution for areas around the 6-bp recognition sequence extending to 2.1 Å (see **Supplementary Figure S3**). This high-resolution data allowed for unambiguous identification and positioning of 25 base pairs of DNA and clear side chain modelling of BrxX core residues (**Figure 2C**), whereas the remaining DNA bases and peripheral loops in BrxX were observed at lower resolution (transparent contour in **Figure 2C**; see also **Supplementary Figure S3D**) due to intrinsic local flexibility.

**Figure 2:**
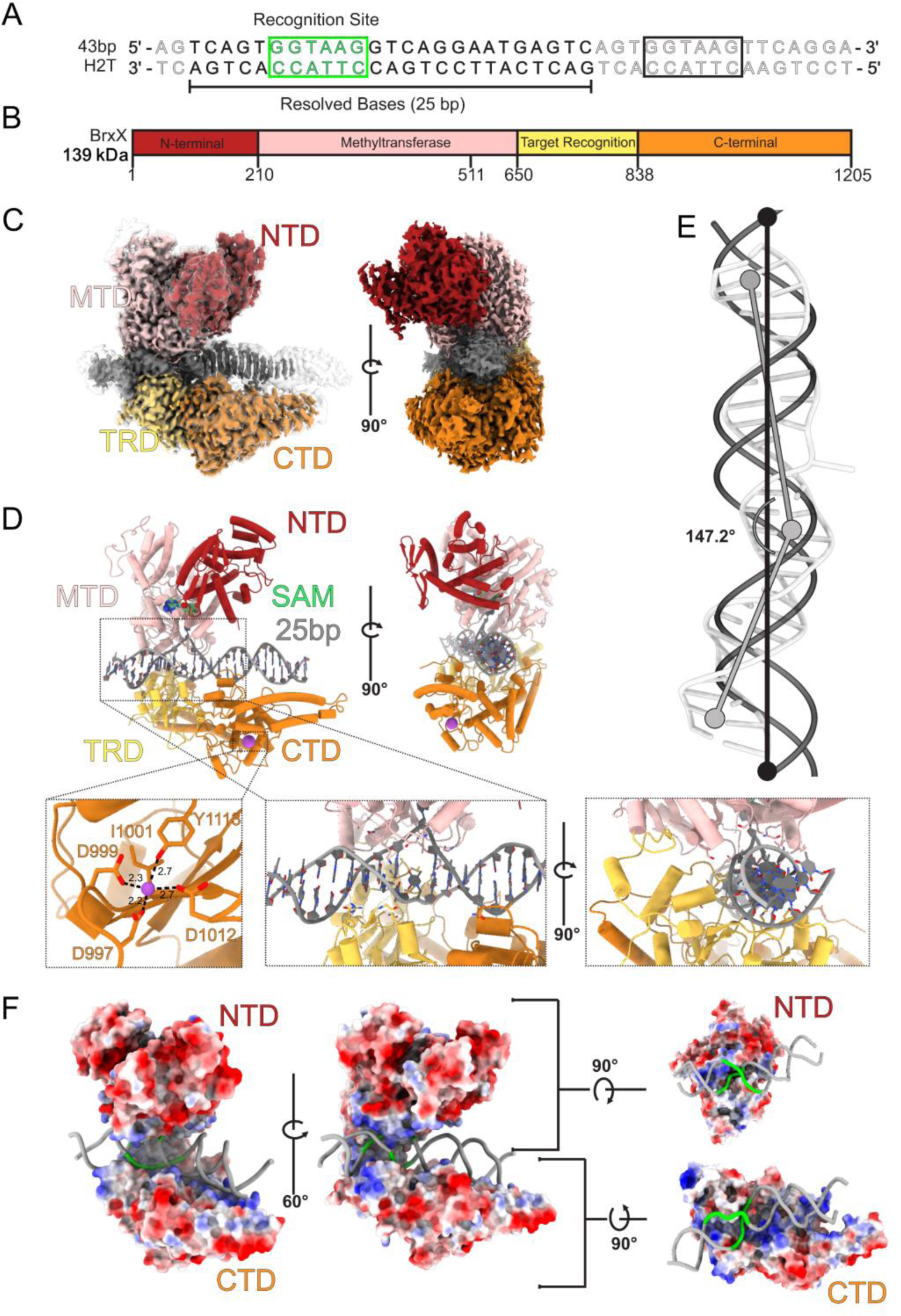
Architecture of BrxX-DNA complex. (**A**) The 43 base pair “Head-to-Tail” (H2T) DNA fragment used in cryoEM sample preparation. The 6 base pair BREX recognition sequence is shown in green, dark grey shows the resolved (modelled) bases whereas light grey shows unresolved bases. Black rectangle highlights the second (unbound) BREX site. (**B**) Domain arrangement of the BrxX monomer: N-terminal domain (NTD) is shown in dark red, methyltransferase domain (MTD) in pink, target recognition domain (TRD) in yellow, and C-terminal domain in orange (CTD). This color scheme is used throughout the manuscript. (**C**) CryoEM maps of the BrxX-DNA complex. The sharpened map, contoured at 5σ, is colored according as described above and shown within the contour of unsharpened map (also at 5σ, transparent contour). Only the density within the sharpened map was modelled. (**D**) A cartoon representation of an atomic model of the BrxX-DNA complex. One SAM molecule is shown as green van der Waals sphere representation. Boxed regions are as follows, L to R: (i) a metal-binding site in CTD and (ii) a DNA-binding region. (**E**). A comparison of the modeled 25 bp BrxX-bound DNA (gray tubes) and ideal B-form DNA (black cartoon). BrxX-bound DNA is bent by ∼33°, and major and minor grooves are stretched. (**F**) Surface coloring based on Coulombic electrostatic potential calculated by ChimeraX^52^ where red is negative potential and blue is positive potential.

DNA-bound BrxX is a bi-lobed “Pacman”-like monomer consisting of four domains. The upper Pacman ‘jaw’ contains the N-terminal domain (**Figure 2BD**, NTD) and the methyltransferase domain (MTD) while the lower ‘jaw’ comprises the target recognition domain (TRD) followed by the C-terminal domain (CTD). The two ‘jaws’ are connected by flexible hinge loops (**Figure 2D**, residues 638-675) at the interface between the MTD and TRD. The NTD comprises an N-terminal α-helix followed by a three-stranded antiparallel β-sheet and a cluster of eight α-helices. The MTD displays a Rossman-like methyltransferase fold comprised of a nine-stranded β-sheet flanked by ten α-helices, and a type IV (NPPY) motif^53^ catalytic pocket with bound SAM. The TRD is responsible for the site-specific DNA recognition and is folded into two α/β subdomains. The CTD consists of the α-helical part and a three-stranded antiparallel β-sheet, connected to a two-helix bundle (the lowermost part of Pacman’s ‘jaw’) by what appears to be a metal-binding site (**Figure 2D**). A metal ion is coordinated by five residues (side chains of Y1113 and N1012, N997, N999) and the main chain oxygen of I1001. As the EM buffer contained Mg^2+^, and because of the octahedral shape of the metal density and six-valence coordination, we hypothesize that the metal ion might be Mg^2+^, but its exact nature remains to be determined.

Conservation analysis of BrxX proteins closely related to *E. coli* HS BrxX by ConSurf server^54^ identified the TRD as the most variable part of the protein, presumably reflecting diversity of sequences recognized by different BREX systems. In contrast, sequences of CTD, particularly around the metal-binding site, and of MTD and NTD, particularly around the SAM binding pocket, were highly conserved (**Supplementary Figure S4A)**. Yet, the NTD and CTD are not well conserved beyond Type I BREX systems (we failed to detect significant structural homology using DALI^55^ or Foldseek^56^ servers), suggesting these domains are unique for Type I systems and have specific recognition or mechanistic functions within a larger BREX complex(es) (see below). The BrxX MTD can be compared to corresponding domains of other DNA methyltransferases (**Supplementary Figure S4B, Supplementary Table S4**): MmeI^57^ and DrdV^29^ (Type IIL R-M enzymes), CamA^58^ (an orphan methyltransferase), and the HsdM subunit of the Type I R-M system EcoKI^59^. BrxX shares highly conserved SAM-binding (motif I) and methylation (motif IV) motifs with these proteins^53^ (**Supplementary Figure S4B**). However, the BrxX MTD contains an insert between residues 390 and 500 (**Supplementary Figure S4C**), which is not observed in other homologs and, alongside with CTD and NTD, could participate in interactions with other BREX proteins.

The DNA bound to BrxX is bent by ∼33° (**Figure 2E**), with the most prominent distortion occurring around the 6-bp recognition sequence, which is embedded between the MTD and TRD. The fifth adenine of the **GGTA<u>A</u>G** sequence is rotated out of the DNA backbone together with its deoxyribose moiety and the neighbouring phosphodiester groups and is nestled in the active site of the MTD. There is a significant rotation of the opposite thymidine as well, and shifting of the surrounding bases resulting in discontinuity of the non-target strand bases π-stacking. The major groove over the recognition sequence widens to ∼25 Å while the minor groove widens to ∼17 Å compared to 19 and 12 Å, respectively, in the B-form DNA. There are a total of 35 direct interactions between the BrxX monomer and DNA; 29 of them occur with the six bases involved in site-specific recognition whereas the 3′ ‘tail’ of the DNA fragment is stabilized by electrostatic interactions with a positively charged groove of the CTD (**Figure 2F**). The distorted state of DNA required for site-specific recognition is stabilized by a series of positively charged residues forming direct contacts with the phosphate backbone. A positively charged lining of the “upper jaw” consists of K226, K522, K515 and K533 of MTD (**Figure 2F** and **Figure 3A**); the complementary positively charged groove in the ‘bottom jaw” includes K719 and R944 of the TRD, and R1145 and K1152 of the CTD that stabilize the bent ‘tail’ of the modelled DNA segment. The overall shape and mode of DNA binding resembles the one observed in MmeI:DNA complexes (**Supplementary Figure S4D**).

**Figure 3:**
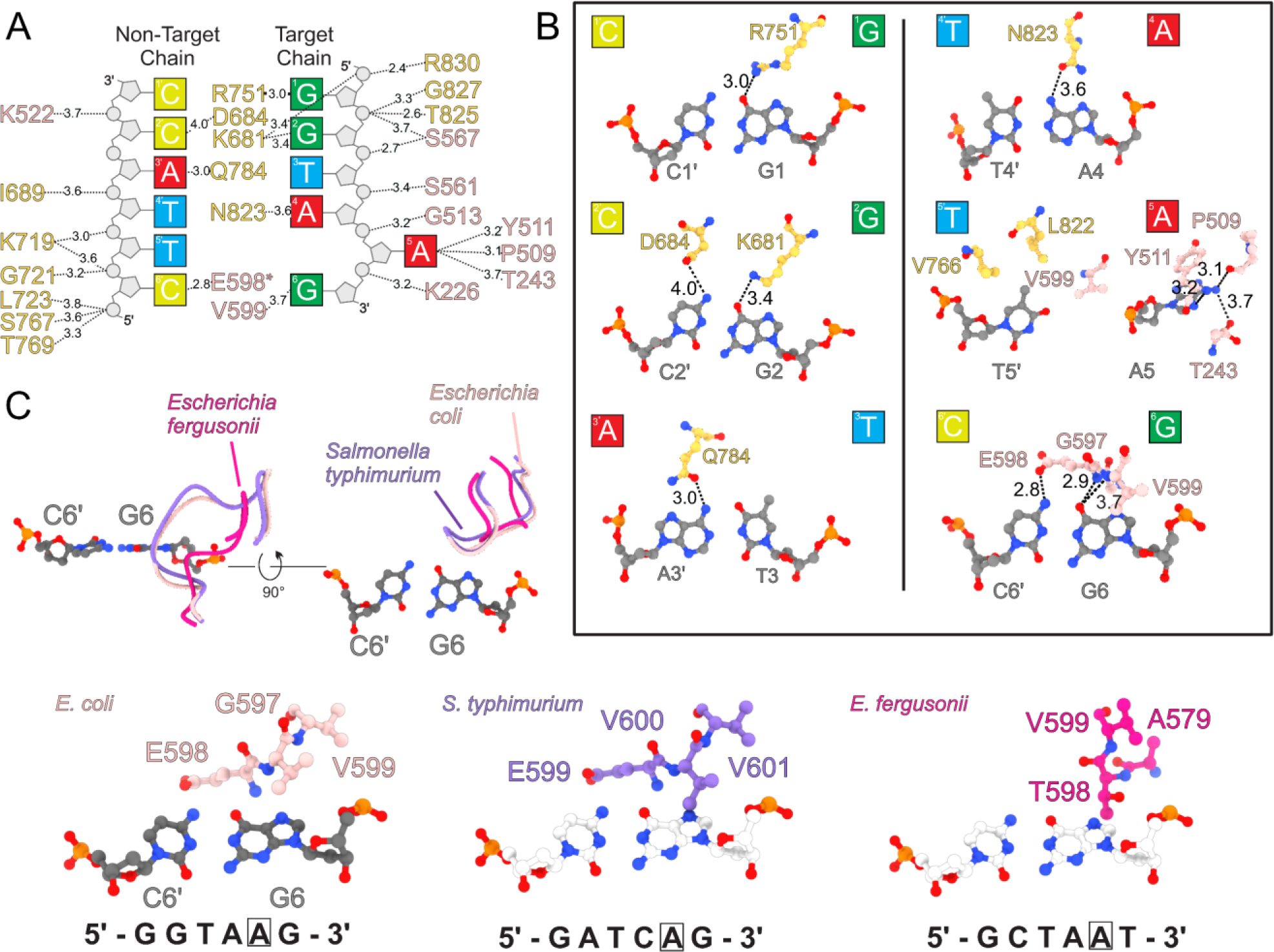
Molecular basis of BREX site recognition by BrxX. (**A**) A scheme of all interactions between BrxX and DNA. Distances in Angstrom are indicated. An asterisk indicates that the main chain and not the side chain is involved. Colors correspond to amino acids from MTD (pink) or TRD (yellow). (**B**) Molecular basis for site-specific recognition of the BREX site. Each base pair is presented separately and oriented with major groove on top. Distances in Angstrom are indicated. (**C**) A species-specific MTD loop recognises the sixth base pair. A structural superposition is shown of loop 587-601 of BrxX MTD and equivalent loops in *Escherichia fergusonii* (based on AlphaFold DB models AF-A0A5A8QD96-F1 and AF-B7L3T0-F1 respectively).

### Structural basis of site-specific DNA recognition by BrxX

All site-specific direct interactions with DNA bases happen within the major groove of DNA to allow base discrimination; they include both the target and non-target DNA strands (**Figure 3AB**). The TRD contributes to the recognition of the first four base pairs of the recognition sequence. Specifically, R751, K681, and N823 recognize G1, G2, and A4 of the target sequence whereas D684 and Q784 recognize C2′ and A3′of the non-target strand (**Figure 3AB**). The flipped adenine is flanked by an A:T base pair which facilitates local DNA buckling and bending that accompanies base flipping. Involvement of residues from the MTD in DNA recognition is not unusual among DNA methyltransferases^29,57,60^, and in our structure occurs with the fifth and sixth positions of the BREX site. Adenine base-flipping is facilitated by the intercalation of hydrophobic residues V766 and L822, which causes a 90° propeller-twist rotation of the opposite T5′ (see **Figure 3AB**), and V599 that sterically fills the empty space that is formed by the appearance of the extrahelical adenine. Together, these three residues create a hydrophobic patch to accommodate the C7 methyl group of T5′. Similar to other MTases, the flipped adenine is inserted deep into the methylation active site (see **Figure 2D**; the adenine ring is stacked against Y511 and surrounded by hydrophobic residues L245, V557, F592, and I595. An amino group of the adenine forms direct interactions with T243 and N508 (3.6 Å and 4.5 Å respectively), facilitating specific base recognition.

Our analysis identified a critical loop (residues 587 to 601) in *E. coli* BrxX MTD that confers specificity to the recognition of the final base pair of the BREX site. Not only does residue 598 make direct contacts with the target strand terminal base G6 (main-chain amide to O6, 2.9 Å) and the opposite C6’ (side-chain oxygen to N4, 2.8 Å; see **Figure 3C**), but the loop sterically enables only the R:Y pairing in this position. The importance of these observation is revealed by comparison with the BrxX homologs from *Salmonella typhimurium* and *Escherichia fergusonii*. These BREX systems recognize, correspondingly, the **GATC<u>A</u>G** and **GCTA<u>A</u>T** sequences^31,32,61^. An AlphaFold2 model of *S. typhimurium* BrxX reveals that the predicted cognate loop around the sixth position of the recognition sequence is strikingly similar to that of *E. coli* BrxX, which is expected given the identity of the terminal base. In contrast, in *E. fergusonii* BrxX this loop is markedly shorter, and is thus able to accommodate the T:A base pair (see **Figure 3C**).

### Ocr sequesters two BrxX monomers and prevents DNA binding

Phage T7 DNA mimic Ocr is an inhibitor and trigger of various immune systems^44,62,63^. In a previous study, we showed that Ocr inhibits BREX defense by binding BrxX *in vivo*^43^. To understand the mechanism of this inhibition we studied BrxX interaction with the Ocr dimer *in vitro*. We incubated BrxX with an excess of Ocr and analysed the sample composition by size-exclusion chromatography (SEC). A shift in the SEC peak of the complex as compared to the BrxX monomer indicated formation of an oligomeric species with a molecular weight likely corresponding to a BrxX_2_:Ocr_2_ tetramer (**Figure 4A**).

**Figure 4:**
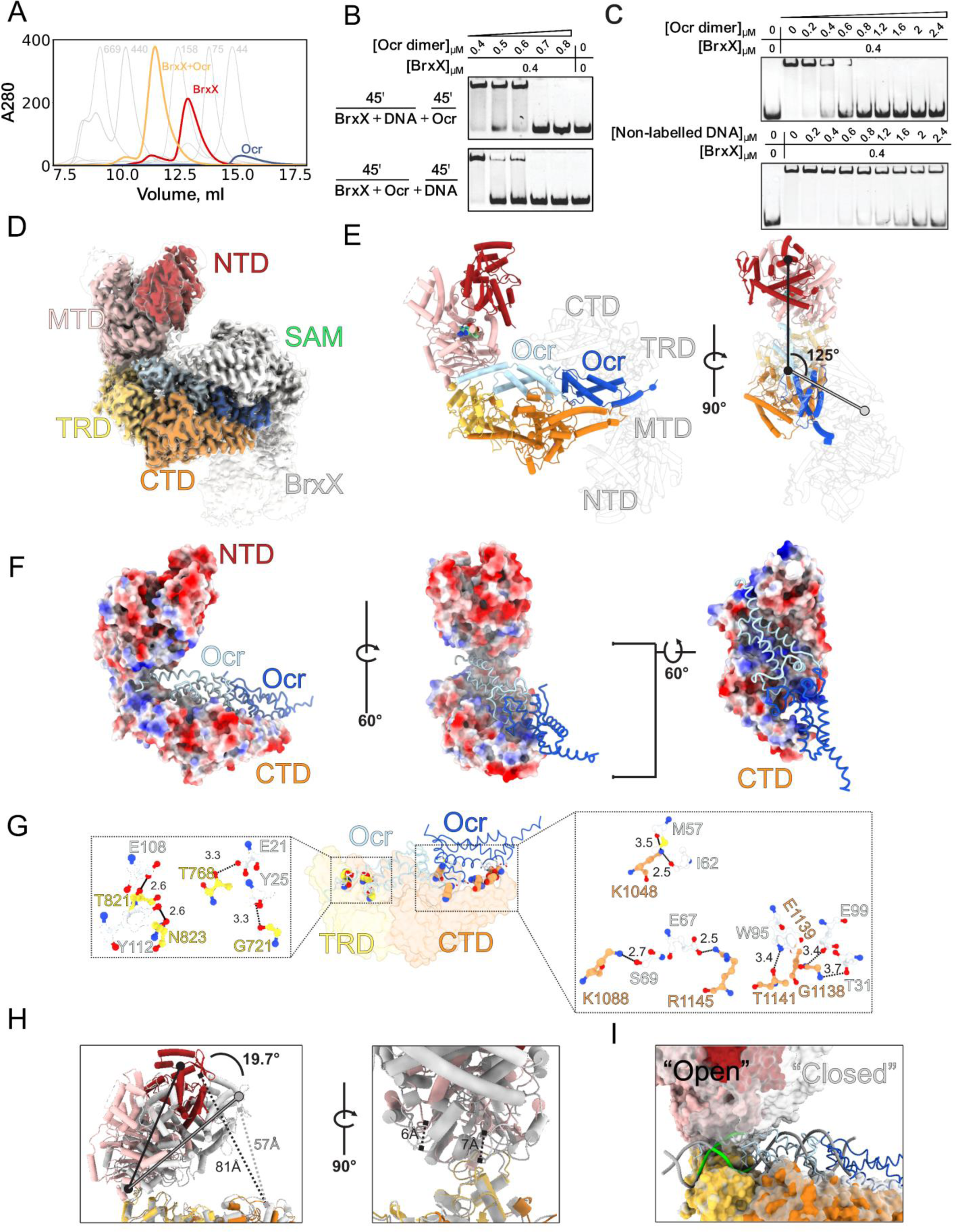
The DNA mimic Ocr displaces DNA from BrxX and locks it in an inactive conformation. (**A**) SEC traces of BrxX, Ocr, or an equimolar BrxX + Ocr mixture injected onto Superdex 200 Increase 10/300 column. SEC profiles of calibrants and their corresponding molecular masses are indicated on the background in grey. (**B**) EMSA with 20 nM Cy5-labelled 40 bp dsDNA substrate with a single BREX site incubated with a 20-fold (0.4 μM) molar excess of BrxX in the presence of indicated concentrations of Ocr dimer. BrxX was either incubated with DNA for 45 min, followed by additional 45 min incubation with Ocr (*top*), or was first incubated with Ocr for 45 min, followed by additional 45 min incubation with DNA (*bottom*). (**C**) Competition between DNA and Ocr for binding to BrxX. 20 nM Cy5-labelled 40 bp dsDNA substrate bearing one BREX site was incubated with a 20-fold (0.4 μM) molar excess of BrxX, followed by an indicated amount of Ocr dimer (*top*) or of a non-labeled dsDNA substrate (*bottom*). (**D**) CryoEM maps of the BrxX-Ocr complex. The autosharpened map, contoured at 7σ, is colored according to BrxX domains coloring scheme (Figure 2) and shown within the transparent contour of the unsharpened map (at 5σ). Ocr monomers are shaded in blue; the lower resolution BrxX monomer is shown in white. (**E**). A cartoon representation of the atomic model of the BrxX-Ocr complex. One SAM molecule is shown as green van der Waals sphere representation. (**F**) Surface coloring based on Coulombic electrostatic potential (as implemented in ChimeraX), where red is negative potential and blue is positive potential. Ocr dimer is shown as worm representation. (**G**) Direct interactions between BrxX and Ocr residues. Interacting residues are shown as balls and sticks; Ocr residues are white while TRD and CTD residues are colored yellow and orange, respectively. (**H**) A structural superposition of atomic models of DNA-bound (white) and Ocr-bound (colored by domain) BrxX monomers. Movements are indicated. (**I**) Structural superposition of surface models. DNA-bound model depicted in transparent white and an Ocr-bound depicted colored by domain. Alignment generated via ChimeraX matchmaker tool using the TRD domain boundaries.

To demonstrate that Ocr competes with DNA for the binding to BrxX, we tested its ability to disrupt the BrxX:DNA complex formation using EMSAs. At our conditions, a 20-fold molar excess of BrxX was sufficient for sequestration of all dsDNA in the reaction. Subsequent incubation with a 1.5-fold molar excess of the Ocr dimer over BrxX completely abolished BrxX:DNA complex formation and released free DNA (**Figure 4B**). Thus, Ocr can efficiently substitute DNA in pre-formed BrxX:DNA complexes. When BrxX was first incubated with Ocr followed by the addition of DNA, an equimolar amount of Ocr dimer was sufficient to block the BrxX:DNA interaction (**Figure 4B**). To directly demonstrate that Ocr is a preferable substrate of BrxX, we compared the ability of Ocr or unlabelled dsDNA to compete with labelled dsDNA for the formation of the BrxX:DNA complex. We found that in both BLI and EMSA Ocr was a better binding partner of BrxX (**Figure 4C, Supplementary Figure S7A-C**).

We next sought to determine the molecular basis of Ocr binding to BrxX by single-particle cryoEM analysis. 2D class averages (**Supplementary Figures S5** and **S6**) showed that the sample contained two distinct particle sets, enabling us to solve both the apo and Ocr-bound structures from a single dataset. 57% of the extracted particles yielded a 2.8 Å global resolution map (**BrxX:Ocr**) corresponding to a BrxX_2_:Ocr_2_ complex where two individual BrxX monomers interact with a dimer of Ocr via their TRDs and CTDs. The two BrxX monomers are oriented at an angle of 125° relative to each other (**Figure 4E**); however, the complex displays imperfect symmetry due to local flexibility of the individual components resulting in varied local resolution (**Supplementary Figure 5E**). We resolved one monomer to ∼2.8 Å (**Figure 4D)** clearly defining all structural features except for a flexible loop between residues 395 and 420. The MTD displayed density for a molecule of SAM, which we modelled based on the higher resolution DNA-bound structure. The map resolution for the second BrxX monomer stretched from ∼3.4 Å around the core of the particle to ∼4-6 Å at the periphery (**Supplementary Figure S5E**); hence, we rigid-body fitted the first monomer model into the density of the second monomer (see **Methods**). Around ∼10% of BrxX particles did not bind Ocr in our sample, enabling us to determine the structure of apo BrxX by rigid-body fitting BrxX:DNA model in a 3.9 Å **BrxX_apo_** map (**Supplementary Figure S6 & Methods**). Given that apo BrxX adopts exact same structural conformation as BrxX bound to Ocr (**Supplementary Figure S6AB**), we proceeded with the higher resolution structure for all further analyses.

Ocr is highly negatively charged, and its interaction with BrxX is largely driven by electrostatic contacts with the positively charged groove on the lower half of the BrxX monomer (**Figure 4F**). To form direct contacts, Ocr utilises two pairs of tyrosine and glutamic acid residues that form hydrogen bond with four residues of BrxX TRD: T82, N823, T768, and the main chain carbonyl of G721 (**Figure 4G**). Three out of four of these TRD residues are also directly involved in DNA recognition or binding: N823 directly interacts with the A4 in the target strand of the recognition site, while the main chain amide of G721 interacts with the phosphate backbone of T5’ base in the non-target strand. T768 lies between S767 and T769, which both interact with terminal phosphate of the non-target strand (**Figure 3A**). Ocr is additionally stabilized by the BrxX CTD via a salt bridge between Ocr E67 and BrxX R1145; the latter residue also interacts with the DNA away from the recognition site. Other CTD interactions include hydrogen bonding between a number of residues (**Figure 4G**).

Structural alignment of the TRD domain in BrxX-DNA and BrxX-Ocr complexes results in Ocr and DNA occupying the same horizontal plane, allowing for a comprehensive comparison of DNA and Ocr geometry and key residue interactions (**Figure 4H**). Structural alignments of each of the four BrxX domains between the two conformations show that the NTD rearranges the least, and the CTD rearranges the most, however all RMSD values fall below 2 Å (**Supplementary Table S4**). The superposition of the two structures makes it clear that it is the ‘upper jaw’ of the BrxX monomer that closes around the DNA upon recognition as the MTD moves 19.7° between the open and closed states (**Figure 4H, left**). The MTD loops that stabilize distorted DNA move 6-7Å from the open to closed state (**Figure 4H, right**). The resolved 25 bp DNA fragment has the same width as the Ocr monomer (**Figure 4I**); however, compared to the relatively rigid Ocr, DNA is a more flexible substrate that can bend and deform. While DNA is clearly able to associate with BrxX in the open conformation, stabilization of a flipped adenine is only possible in the closed conformation which brings each strand of the DNA substrate in close contact with BrxX, and facilitates the formation of stabilizing contacts between the MTD and the distorted phosphate backbone of bound DNA.

### BrxX lacks methyltransferase activity *in vitro*

S-adenosyl-L-methionine (SAM) is a universal methyl group donor to C, O, N or S atoms and thus an essential cofactor of methyltransferases^64^. In both DNA-bound and Ocr-bound BrxX structures, a clear density for a SAM molecule is visible deeply nestled within the hydrophobic pocket in the core of the MTD (**Figure 5A**). Apart from the narrow channel leading from the flipped base pocket to the active site, the SAM molecule is otherwise completely surface inaccessible. The lack of significant rearrangement between the apo- and DNA-bound conformations demonstrates that BrxX, at least in the presence of SAM, is in a methylation-primed state, and has a pre-organized binding pocket to receive the flipped base. This observation is in line with the EMSA results showing the enhancing effect of SAM on DNA binding (**Figure 1C**), and with our cryoEM data. SAM was critical to observe high-resolution 2D classes, suggesting that SAM binding limits flexibility and helps to organise the MTD (data not shown). All three regions of the SAM molecule (adenine ring, ribose, methionine moiety) make extensive contacts with BrxX (**Figure 5B**). The adenine ring is stacked between F540 and I355 and further stabilized by VdW contacts with L536 and I239. The ribose ring forms direct hydrogen bonds with Q244, and methionine end of the molecule is also encapsulated within an extensive network of H-bonding (**Figure 5B**). The NPPY motif IV sequence (508-511) accommodates the target adenine flipped out of the DNA helix, with Y511 playing a key role in adenine stabilisation. In line with these observations, the Y511A mutation disrupted both site-specific and non-specific DNA binding (**Figure 5C**), and completely abolished the BREX defense and methylation *in vivo*, without causing cell toxicity (**Figure 5D**). Inflection temperature analysis using Tycho NT.6 confirmed that the Y511A mutation did not affect the overall stability of the protein (**Supplementary Figure S7D**). These results indicate that anchoring of the MTase domain by a flipped-out adenine following the transition of BrxX to the closed conformation is an essential step in BREX site recognition.

**Figure 5:**
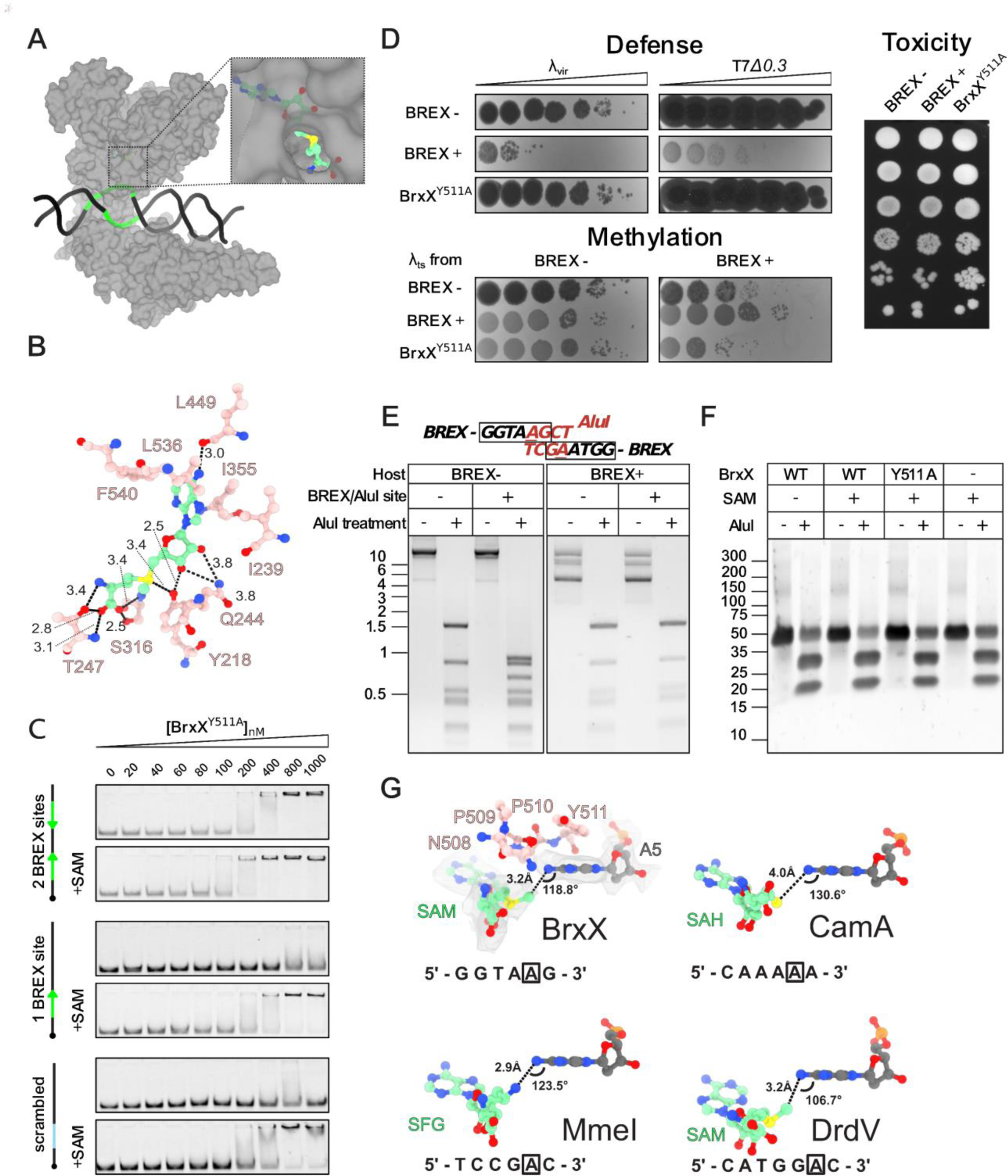
BrxX alone lacks methyltransferase activity. (**A**) Surface representation of the BrxX monomer shown in grey bound to DNA (black) containing a BREX site (green). Inset highlights the only external access point to the SAM molecule enclosed within the catalytic pocket. (**B**) Direct interactions between the SAM molecule and MTD residues. Dashed lines indicate hydrogen bonding with distances indicated in Angstroms. (**C**) Effects of BrxX^Y511A^ mutation on BREX defense, methylation, and toxicity. BREX defense is demonstrated by an EOP assay with BREX-sensitive *λvir* and T7Δ*0.3* phages. BREX methylation is estimated by ability of *λts* induced from the indicated lysogenic cultures to plaque on BREX^-^ and BREX^+^ lawns (see **Methods**). Toxicity was estimated in a drop-spot test on LB agar plates. BrxX^Y511A^ was introduced in the context of the full BREX cluster. All assays were performed in biological triplicates and representative plates are shown. (**D**) EMSA with 20 nM Cy5-labelled 43 bp dsDNA substrate without BREX sites (“scrambled”), or with one or two BREX sites, incubated with indicated amount of BrxX^Y511A^ without co-factors or in the presence of 0.5 mM SAM. Green arrows indicates orientation of BREX sites. Compare with Figure 1C for EMSA with wild-type BrxX performed in identical conditions. (**E**) Overlapping BREX methylation inhibits AluI cleavage. An agarose gel shows AluI digest of pHERD30t or pHERD30t bearing an overlapping BREX/AluI cleavage site, purified from either BREX^+^ (*E. coli* HS) or BREX^-^ (*E. coli* BW25113) culture. (**F**) AluI cleavage with 40 bp dsDNA substrate containing an overlapping BREX/AluI site and incubated with 100 μM BrxX overnight. (**G**) A comparison of orientations of the flipped adenine and SAM or an equivalent ligand between BrxX (this work), CamA (PDB: 7LT5), MmeI (PDB: 5HR4), and DrdV (PDB: 7LO5). The 5’ to 3’ recognition site with the methylated adenine boxed is indicated for each protein. In the BrxX structure, the sharpened map density contoured at 5σ is shown surrounding each molecule, with the catalytic residues (NPPY motif) from the MTD shown in pink.

To detect BrxX methyltransferase activity *in vitro* we relied on the fact that the AluI restriction endonuclease is sensitive to adenine methylation^65^, and that the AluI recognition site (**<u>A</u>GCT**) overlaps with the BREX site (**GGTA<u>A</u>G**). To demonstrate AluI sensitivity to BREX methylation we constructed a plasmid carrying an overlapping BREX/AluI site and confirmed that when purified from BREX^+^ *E. coli* HS cells, this site is not cleaved by AluI in contrast to the same plasmid purified from BREX^-^ cells (**Figure 5E**). Next, we used a 40 bp dsDNA substrate carrying a single BREX/AluI and a single Dam site in the *in vitro* methylation reaction with wild-type BrxX or BrxX^Y511A^ mutant. Dam methyltransferase was used as a positive control. While Dam methylation was complete after overnight incubation with the protein, as revealed by the sensitivity of methylated DNA to DpnI/II cleavage (**Supplementary Figure S7E**), substrate incubated with BrxX was efficiently cleaved by AluI, indicating the lack of methylation (**Figure 5F**). To exclude the possibility that in our conditions BrxX methylates its target sites at a slow rate that is beyond the detection limit of a restriction-sensitivity assay, we attempted to detect N6-methyl-adenine (m6dA) using HPLC-MS analysis of DNA substrate digested to nucleosides (**Supplementary Figure S7F**). Again, while the amplitude of the m6dA signal in Dam-treated sample corresponded to full methylation of a single Dam site of the substrate, no signal was detected in BrxX-treated sample, confirming that BrxX alone lacks methyltransferase activity using the conditions of our *in vitro* assay.

In the DNA-bound BrxX structure, the N6 amino group of the extrahelical adenine forms hydrogen bonds with the side chain of N508 (4.3 Å) and the backbone carbonyl of T243 (3.6 Å). This would negatively polarize the N6 atom and activate it for a direct transfer of the methyl group from SAM via nucleophilic substitution using Sn2 proximity and desolvation mechanism. However, for this to occur, the three atoms (N, S C) must form a linear arrangement. As the two catalytic moieties are at an angle of 118.8° relative to one another, methylation in this state would not be possible indicating a pre-, or post-methylation state. This is no different from other similar methyltransferases such as MmeI^57^, CamA^58^, and DrdV^29^, where the structures show a similar distance and angle orientation of the N6 adenine atom and the methyl group carbon atom (or the equivalent moiety in the respective structures) (**Figure 5G**). None of these structures directly show how the methylation reaction occurs, but as all these enzymes methylate their substrates *in vitro*, we conclude that the elements present only in BrxX (such as NTD, CTD, or the MTD insert) might act as cis-inhibitory elements preventing BrxX from occupying a transient methylation-competent conformation, and that interactions with external factors are required to relieve this inhibition.

### BREX methylation and defense require assembly of a macromolecular BrxBCXZ complex

To identify conditions supporting BREX methylation *in vivo*, we expressed BrxX (alone, or in combination with other BREX proteins) in *E. coli* cells lysogenized with the *λ* phage. Induction of a prophage from BREX^+^ cells resulted in the production of methylated phage progeny that bypassed BREX, while the non-methylated phage produced by the BREX^-^ culture was restricted on a BREX^+^ lawn^18^ (**Figure 6A**). Phage induced from cells expressing BrxX was fully susceptible to BREX defense, indicating the lack of BREX methylation (**Figure 6A**) and in line with *in vitro* results (**Figure 5EF**). Co-overproduction of BrxX with any other single BREX protein also did not support *in vivo* methylation (data not shown). However, phages induced from cells simultaneously producing BrxB, BrxC, BrxX, and BrxZ proteins were resistant to BREX (**Figure 6A**), consistent with our previous observation that individual deletion of each of these components abolished BREX methylation^18^. This result shows that methylation-competent state of BrxX requires BrxB, BrxC, and BrxZ proteins, and suggests that the four proteins may physically interact.

**Figure 6:**
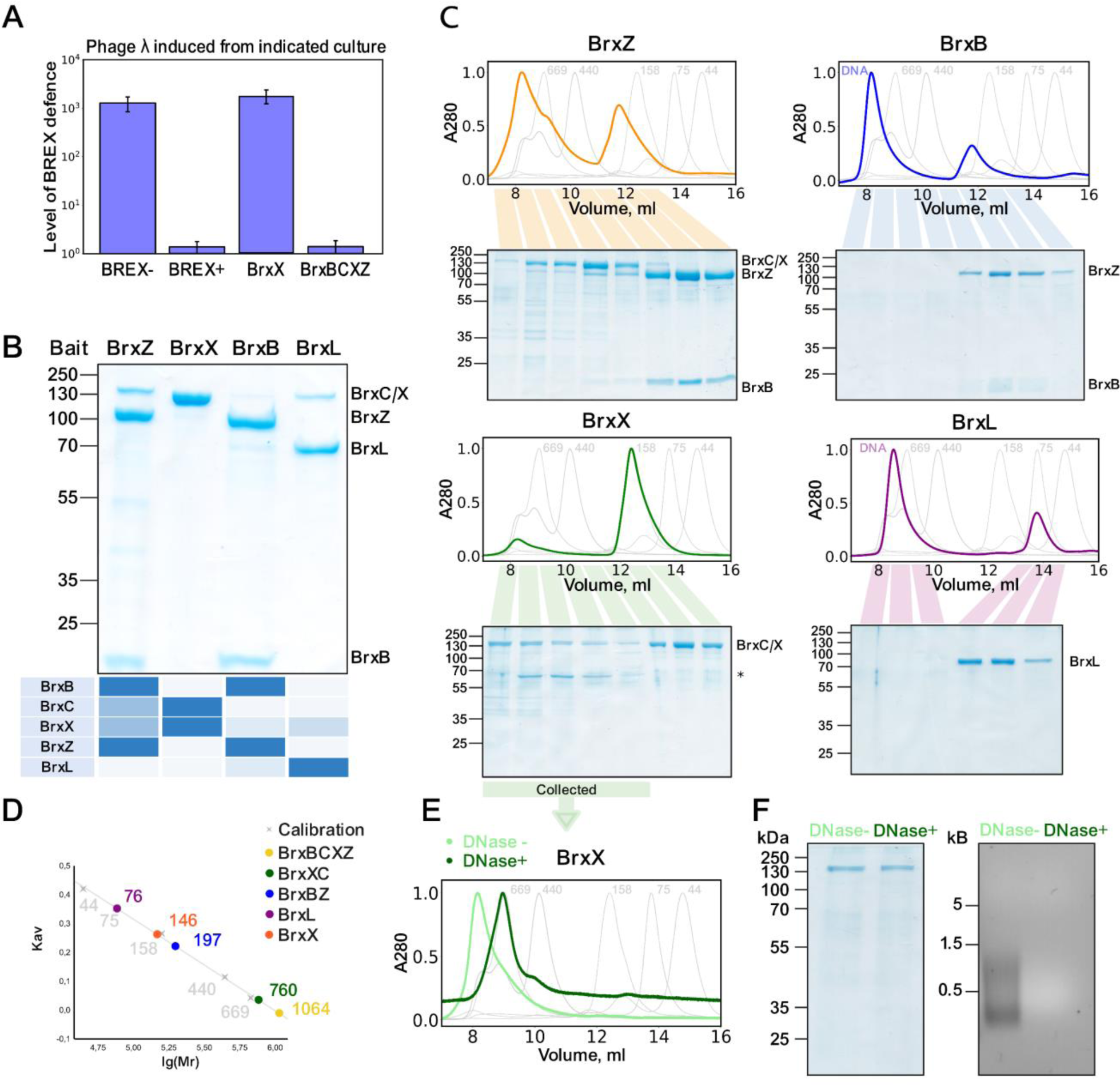
BREX methylation requires assembly of a macromolecular BrxBCXZ complex. (**A**) *in vivo* BREX methylation requires co-production of BrxX, BrxC, BrxZ, and BrxB. Prophage *λwt* was induced from indicated lysogens and the status of its BREX methylation was determined by plaquing on BREX^+^ and BREX^-^ lawns. (**B**) *In vivo* pull-downs with Strep-tagged BREX proteins expressed in the context of the full BREX cluster. Strep-tagged protein (baits) are indicated at the top and co-eluted proteins, identified through MALDI-TOF mass-spectrometry, are shown on the heatmap below. The Strep-trap column eluates were concentrated, and proteins resolved by 4-20% gradient SDS-PAGE. (**С**) Strep-tag pull-down eluates (**B**) were concentrated and separated on a Superdex 200 Increase 10/300 column. For each run, 280 nm absorbance was normalized to the highest value. SEC profiles of calibrants and their corresponding molecular masses are indicated on the background in grey. An asterisk indicates contaminant protein. DNA non-specifically binds Strep-Trap HP column, creating peaks eluting with the void volume, which are the most evident in BrxL and BrxB runs. (**D**) SEC calibration curve with estimated masses of obtained protein complexes. (**E**) DNAse I treatment of fractions containing the BrxCX complex results in the reduction of apparent complex size. (**F**) SDS-PAGE results (*left*) demonstrating protein and agarose gel electrophoreses (*right*) demonstrating DNA content of the BrxCX peak before and after DNase I treatment. In all panels, the protein or DNA molecular weight markers are shown on the left.

To systematically determine protein-protein interactions of BREX components we introduced a sequence encoding Strep-tag II onto the 3’ end of each *brx* gene individually within the context of a full BREX gene cluster. Strep-tagged BrxA and BrxC variants did not show phage defense (**Supplementary Figure S7G**). We therefore carried pull-down assays only with Strep-tagged BrxB, BrxX, BrxZ, and BrxL constructs (**Figure 6B**). The results support the existence of a BrxBCXZ complex, which is detected when BrxZ-Strep is used as a purification bait. SEC analysis of BrxZ-Strep eluate showed two separate peaks corresponding to BrxZB and BrxBCXZ complexes (**Figure 6C,D**). The BrxBCXZ complex co-purified with cellular DNA and thus migrates close to the void volume of a gel filtration column, preventing correct estimation of molecular weight and stoichiometry of the complex. Pull-down with BrxB confirmed that this protein directly binds to BrxZ, resulting in a formation of a stable ∼200 kDa complex (**Figure 6CD**). BrxX-Strep co-purified with BrxC, and also with the host DNA, however, DNase I treatment allowed us to estimate the molecular weight of BrxCX as ∼760 kDa, indicating formation of oligomers (**Figure 6C-F**). Compared to other tagged BREX proteins, the level of BrxL-Strep production was the lowest and only purification from a larger volume of culture allowed us to detect its interaction with BrxX (**Figure 6CD**).

Together, these results support a structural model for the assembly of a BREX complex where BrxB preferentially interacts with BrxZ, and BrxX - with BrxC. Given that all four proteins are required for both BREX defense and methylation^18^, a single BrxBCXZ complex likely combines both activities and relies on BrxX for DNA recognition, methylation of host DNA, and sensing non-methylated DNA of invading mobile elements. We hypothesize that conserved N- and C-terminal domains of BrxX are required for the interactions with BrxZ and BrxC proteins, and that these interactions alter the conformation of BrxX to make it methylation-competent. How BREX system coordinates defense and methylation activities within a single complex and whether it experiences structural re-arrangements in the process of invading DNA exclusion represents an important question for further studies.

## DISCUSSION

BREX is a representative of a cohort of immunity systems that encode DNA modification components yet lack obvious endonucleases that destroy unmodified DNA in R-M systems. BREX relies on BrxX (PglX) methyltransferase to epigenetically modify host DNA and discriminate it from unmodified DNA of genetic invaders. The mechanism of subsequent BREX defense, which limits the propagation of invaders with unmodified DNA, is yet to be determined. Here, we show that BrxX is the only BREX component responsible for the recognition of BREX sites, and thus is involved in both the sensing of viral non-methylated DNA and the initiation of the BREX defense. The dual role of BrxX is confirmed by an observation that a mutation in the methylase active site (Y511A) inactivates both methylation and BREX defense, but does not result in toxicity, as would be expected if another BREX protein was directly involved in defense upon the recognition of non-methylated host DNA. This result aligns with results of scanning mutagenesis of *Salmonella enterica* BrxX (PglX), which resulted in multiple non-toxic mutants that retained the *in vivo* methylation activity, while reducing BREX defense^34^. On its own, BrxX does not demonstrate methyltransferase activity either *in vitro* or *in vivo*. *In vivo*, co-expression of BrxX with BrxB, BrxC, and BrxZ is required for BREX methylation. We show that these four proteins form a complex, which likely is a minimal unit required for methylation and BREX defense.

Combining both self/non-self recognition and defense functions in one complex is highly advantageous for an immune system that methylates only one DNA strand within an asymmetric recognition motif. Since non-methylated BREX sites inevitably emerge in host DNA after each round of replication, BREX defense must be tightly regulated to avoid potentially lethal activation by non-methylated sites (**Figure 7A**). The principal difference between host and phage DNA is that the latter is completely non-methylated when entering the host cell, while replicated host DNA contains non-methylated DNA sites only on the non-template strand (**Figure 7A**). Thus, discrimination between self and non-self DNA likely requires an assessment of the methylation status of two neighbouring BREX sites located on different DNA strands. Type III and Type ISP R-M systems, which also methylate only one DNA strand of non-palindromic DNA motifs, rely on such a strategy and recognize two non-methylated sites in Head-to-Head orientation to activate DNA cleavage^24,28^.

**Figure 7:**
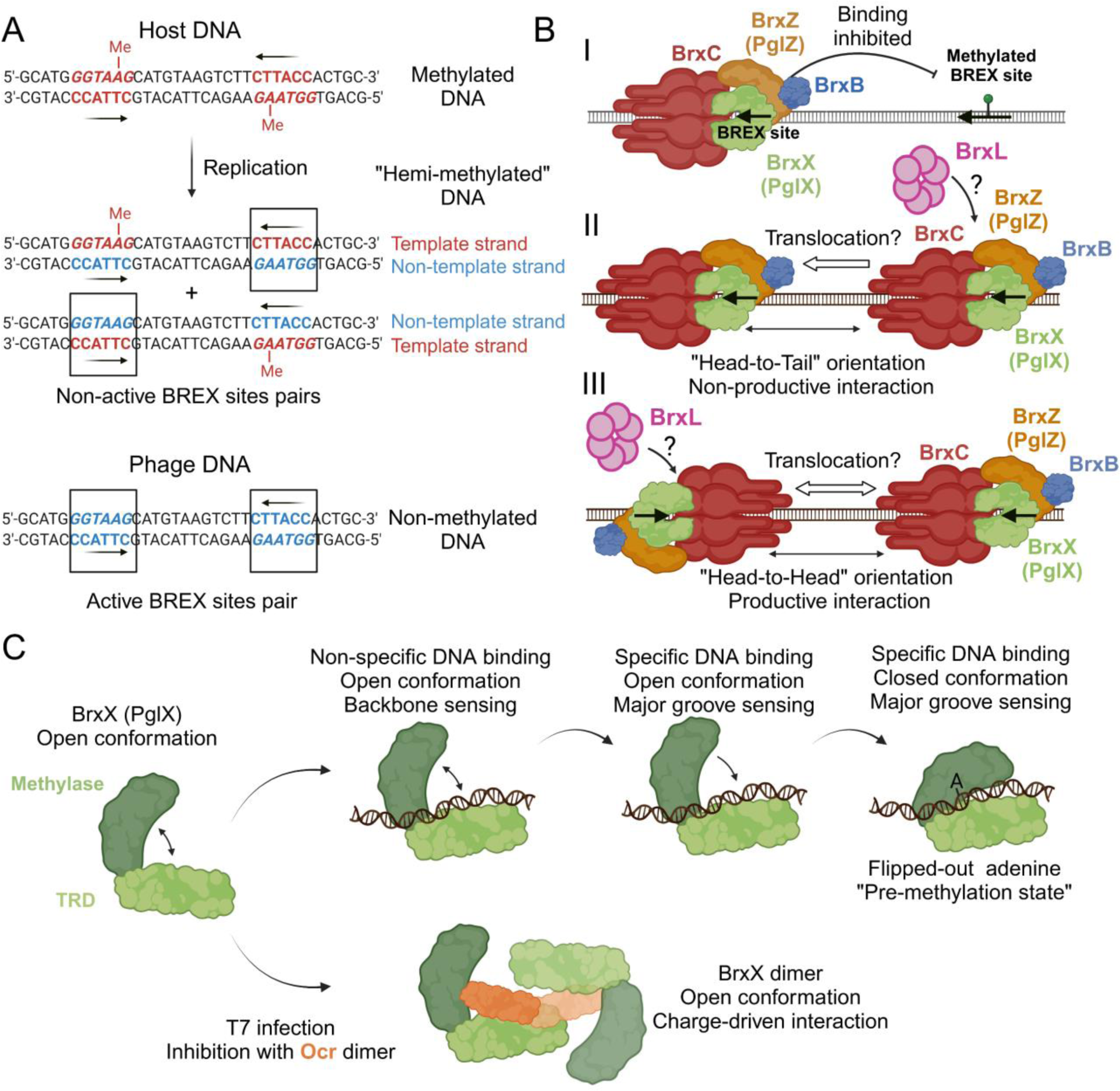
A hypothetical model of BREX defense and invading DNA recognition. (**A**) Replication of host DNA results in generation of non-methylated BREX sites on the non-template DNA strand. (**B**) Discrimination between non-methylated BREX sites in the host and phage DNA requires assessment of the methylation status of two neighboring sites in Head-to-Head orientation. (**C**) BREX sites recognition requires target adenine flipping and BrxX conformation transition from an open to the closed state, while Ocr locks two BrxX monomers in an open conformation.

A hypothetical model for BREX defense is presented in **Figure 7A**. BrxX is a sensor of BREX sites’ methylation within a multifunctional BrxBCXZ complex. Through BrxX, the BrxBCXZ complex recognizes and binds non-methylated BREX sites. Methylation inhibits the binding (**Figure 7B, Stage I**). Binding of BrxBCXZ to two non-modified BREX sites located on different strands (i.e., in a Head-to-Head orientation), identifies DNA as non-self after the two BrxBCXZ complexes interact with each other. Binding to sites in a Head-to-Tail orientation (mimicking the orientation of BREX sites in the non-template strand of replicated host DNA) does not lead to productive interaction, that is, does not activate the BREX defense (**Figure 7B, Stage II/Stage III**). An interaction between the BrxBCXZ complexes likely requires DNA translocation. BrxC is a member of AAA ATPase family, and shares homology with ORC/Cdc6 (Origin Recognition Complex) proteins involved in initiation of archaeal and eukaryotic DNA replication^36,37^. BrxC may play a role of the assembling scaffold/recruitment unit for the complex, similar to the role of ORC, which loads the replicative helicase MCM onto replication origins^66,67^. Strikingly, BrxL protein shares a distinct homology with the MCM helicases^38^ and thus could represent a tentative BREX translocase. Supporting this hypothesis, Type II, III, V, and VI BREX systems that lack BrxL encode BrxH helicase instead. BrxL is not required for DNA methylation but was shown to be required for BREX defense^38^. Another subunit of the BREX complex, BrxZ (PglZ), is homologous to the effector of the PorXY two-component system^35^. PglZ domain of PorX is a phosphodiesterase cleaving cyclic and linear oligonucleotides^35^ and analogous function within BrxZ can be responsible for the effector response to phage infection.

Interestingly, we show that BREX methylation also strictly requires the components of the BrxBCXZ complex and does not proceed with BrxX alone, despite the presence of both adenine and SAM in the methyltransferase domain catalytic pocket of DNA-bound BrxX. Organisation of the BrxX catalytic pocket in the DNA-bound structure is similar to those observed in the structures of other methyltransferases bound with SAM, SAH, or sinefungin^29,57,58^ (**Figure 5G**). These proteins are catalytically active *in vitro* and given their similarity with BrxX, lacking *in vitro* methyltransferase activity, we suggest that the latter structures likely capture either a “pre-methylation”, or a structurally similar “post-methylation” conformation. We speculate that BrxX CTD or NTD domains inhibit transition to a methylation-competent state, and only upon the interaction with other BREX proteins does the conformation change to license methylation of target DNA. Some evidence to support this notion comes from the study of PglX from the Pgl phage immunity system of *Streptomyces coelicolor* A3(2) that lacks NTD and CTD homologous to BrxX and actively methylates DNA *in vitro*^23^.

Comparisons of the apo-form, Ocr- and DNA-bound BrxX structures provide additional insight into events preceding the methylation reaction (**Figure 7C**). TRD and methyltransferase domains of BrxX are connected by a flexible linker, and the apo-protein exists in an ‘open’ conformation. BrxX is able to bind DNA non-specifically, likely through electrostatic interactions of DNA phosphodiester backbone with the TRD and CTD. The Ocr DNA mimic protein exploits this property of non-specific DNA recognition to sequester BrxX. Thus, Ocr interaction with BrxX homologs should not depend on the recognition site of a particular system, explaining its broad inhibitory activity. Following a non-specific DNA binding event, BrxX might scan neighbouring DNA in search of the BREX site, recognition of which is mediated through direct sensing of bases in the DNA major groove and the establishment of an extensive network of additional TRD-DNA contacts. The flipping of the fifth adenine upon the ‘closure’ of the flexible methyltransferase domain is concomitant with the recognition of the last, sixth base by the closed MTD resulting in a stable “pre-methylation” complex. Substitution of BrxX Y511, which forms a π-stacking contact with the extrahelical adenine, results in a loss of both DNA binding *in vitro* and BREX defense *in vivo*. Thus, the “pre-methylation” complex is required for stabilization of the BrxX:DNA interaction and productive BREX site recognition.

Stabilization of the BrxX:DNA complex also explains the role of SAM in BREX defense. In Type I R-M systems, SAM serves not only as a donor of methyl groups but is also required as a co-factor of the restriction reaction^45^. We show that the BrxX:DNA interaction is SAM-dependent: SAM enhances non-specific DNA binding yet has a more prominent effect on BrxX binding to BREX sites. This could be explained through an overall stabilization of BrxX methyltrasferase domain in the presence of SAM, and an improved ability to accommodate flipped out adenine in the catalytic pocket. Since BrxX is involved in sensing of non-methylated sites in phage DNA, SAM depletion should decrease the ability to recognize foreign DNA, rendering BREX defense inefficient. This explains our previous observation of BREX defense inhibition by the phage T3 SAM lyase^46^ and highlights the role of these enzymes as efficient inhibitors of SAM-dependent immunity systems.

To summarise, in this work we have described the molecular basis of DNA recognition by BREX anti-phage defense system, the first step in BREX-mediated immunity. Our results show that in addition to the methyltransferase function BrxX is a sensor of phage infection. We have also revealed how anti-defense protein Ocr sequesters and inactivates BrxX. Finally, we have demonstrated that both BREX defense and methylation require a supramolecular BrxBCXZ complex. Combination of methylation and defense functions in one complex has implications for the evolution and engineering of BREX specificity. In contrast to Type II R-M systems, which are evolutionary constrained by the necessity to change target specificity through accumulation of mutations in both the R and M components but similar to Type I, III and IIL systems^57,68^, novel BREX specificity could emerge upon a single mutation in the *brxX* gene. This implies that BREX could more efficiently counteract accumulation of escaper phages through changes of its recognition motif and might explain previously observed instances of duplications or phase variation of *pglX* gene in BREX clusters^17^. Our DNA-bound structure provides the molecular basis for BREX DNA recognition and can be used as a blueprint for design of BREX variants with desired target specificity.

## Acknowledgements

Skoltech team was supported by RSF grants (22-14-00004 and 22-74-00126). We thank Olga Musharova, and Ekaterina Savitskaya who sadly passed away, for the help with Strep-Seq sequencing, Alina Demkina for the help with plasmids sequencing, and Evgeny Klimuk for the help with protein purification. D.G. is a recipient of a Sir Henry Dale Fellowship (221868/Z/20/Z) funded jointly by the Royal Society and Wellcome Trust; work in his lab is also supported by the BBSRC-funded Institute Strategic Programme “Harnessing Biosynthesis for Sustainable Food and Health” (HBio) (grant number BB/X01097X/1). CryoEM data were collected in the Astbury Biostructure Laboratory (ABSL) CryoEM facility; we thank all staff, particularly Yehuda Halfon for excellent technical support. 200 kV test cryoEM data were collected at John Innes Centre BioImaging facility with the help of Jake Richardson. T.R.B. was supported by the Royal Society International Exchange Grant (IEC\R2\202085), and a Lister Institute Prize Fellowship. K.L.M. is supported by the Canadian Institutes of Health Research (PJT-165936).

## Author contributions

A.I., K.S., and D.G. initiated the study, A.D., M.S., K.P., and A.I. performed biochemistry experiments, M.A. prepared cryoEM samples, collected and analysed cryoEM data. M.A. built and analysed atomic models. M.M. constructed Strep-tagged plasmids and contributed to the *in vivo* pull-downs. A.I., D.G., K.S., and T.R.B. acquired funding. A.I., D.G., K.S., T.R.B., and K.L.M. provided resources for this study. A.I., M.A., A.D., M.S. and D.G. prepared figures and the initial manuscript draft. A.I., D.G., and K.S. edited the manuscript with comments from T.R.B. and K.L.M. All authors have read and agreed with the final version.

## Data availability

All data needed to evaluate the conclusions in the paper are present in the paper and/or the Supplementary Materials, or available upon reasonable request t to the lead contacts – Dr. Artem Isaev and Dr. Dmitry Ghilarov. Sequencing data have been deposited in NCBI database with accession number PRJNA1077651. The atomic models coordinates for BrxX:DNA, BrxX:Ocr and apo BrxX have been submitted to the Protein Data Bank (https://www.rcsb.org/) with PDB IDs 9EWZ, 9EX7 and 9EXH, respectively. Corresponding EM maps have been submitted to the Electron Microscopy Data Bank (https://www.ebi.ac.uk/pdbe/emdb/) with IDs EMD-50027, EMD-50029, EEMD-50032, EMD-50028 and EMD-50038. Raw data were submitted to the Electron Microscopy Public Image Archive (https://www.ebi.ac.uk/pdbe/emdb/empiar/) with IDs EMPIAR-XXX, EMPIAR-YYY.

## Methods

### Bacterial strains and bacteriophages

All bacterial strains, plasmids, and bacteriophages used in this study are listed in **Supplementary Table S1**. The majority of experiments were performed with *E. coli* BW25113. BREX^-^ is a BW25113 carrying an empty pBTB-2 vector, while BREX^+^ carries a full 6-gene *E. coli* HS BREX cluster cloned in pBTB-2 under native promoters^18^. Unless stated otherwise, bacterial cultures were propagated in LB medium (Lysogeny Broth: 10 g/L NaCl, 10 g/L tryptone, 5 g/L yeast extract) at 37°C with appropriate antibiotics. *λ_vir_* is an obligatory lytic mutant of the phage *λ*. *λ_ts_* is a *cI857 bor::Cm* variant encoding thermosensitive CI repressor, allowing for a temperature-controlled lysogen induction. For the infection with phage *λ* cells were supplemented with 0.2% maltose and 5 mM MgSO_4_. T7*Δ0.3* lacks *0.3* gene encoding anti-BREX protein Ocr and thus is sensitive to BREX defense.

### Plasmid construction

Primers used to generate plasmids are listed in **Supplementary Table S2**. pBAD vectors for His-tagged BREX proteins expression and Strep-tag variants of the pBREX AL vector were obtained previously^18,43^. Y511A mutation in pBAD-His-BrxX was introduced using Q5 site-directed mutagenesis kit (NEB), while introduction of Y511A to the pBREX AL was carried by joining 2 plasmid fragments, carrying mutation in the overlap region, using NEBuilder HiFi DNA assembly kit (NEB). pBAD-His_6_-Ocr was obtained by cloning *0.3* gene from T7 phage into pBAD-HisB^43^ vector using Gibson Assembly Master Mix (NEB). Introduction of the overlapping BREX/AluI site to the pHERD30t vector was also achieved via Q5 site-directed mutagenesis kit (NEB). Plasmids were transformed into chemically competent XL1-Blue cells (Evrogen). The presence of desired mutations was verified by Sanger sequencing with specific primers, while absence of secondary mutations was confirmed via whole-plasmid BGI sequencing.

### Monitoring dynamics of phage DNA injection with a potassium efflux assay

Monitoring potassium efflux requires culture infection at high MOI. First, we identified that at MOI of 5 potassium efflux can be robustly detected, while BREX anti-phage defense is still efficient. λ *cI_857_ S_am7_* was used to produce high-titer stock (∼2*10^12^ pfu/ml). A λ *cI_857_ S_am7_* lysogen was grown in 30 mL LB at 30°C for 2 h 30 min, and the prophage was inducted with a 5 min heat shock at 45°C in a water bath. Cells were then incubated at 37°C for 2 h for phage particle production. λ*cI_857_ S_am7_* cannot lyse the membrane, allowing phage particles accumulation inside the cell. Cells were collected by centrifugation (6 000 g, 10 min, 4°C), resuspended in 2 ml of SM3 buffer (10 mM Tris-HCl pH 7.5, 10 mM NaCl, 4 mM MgSO_4_) and lysed by sonication. The cell debris was then removed by centrifuging the lysate for 10 minutes at 6 000 g. The resulting phage lysate was dialysed overnight at 4°C against 1 L of SM3 buffer to remove residual K^+^ and the titer was measured at 37°C on a lawn of the *E. coli* amber suppressor strain QD5003.

For potassium efflux assays, BW25113 cells were grown at 37°C in 15 ml LB supplemented with 0.2 % maltose and 10 mM MgSO_4_ to an OD_600_ of 0.5. A 5 ml aliquot was removed, the cells were collected by centrifugation at 3,000 g for 5 min, and the resulting cell pellet was resuspended in 5 ml of SM3. The centrifugation/resuspension step was repeated three times to completely remove residual K^+^ ions.

The Orion potassium selective electrode (Thermo Scientific) was assembled and stored in 10 mM K^+^ buffer for 24 hours. Before the assay, the electrode was adopted to low concentrations of K^+^ (10^-7^ – 10^-5^ M K^+^ in buffer with Ionic Strength Adjuster). A standard curve of K^+^ ions concentration was built before each experiment in a range of 10^-6^ - 10^-3^ M of K^+^. The electrode was equilibrated with 2 ml of cells for 5 min and then 100 μl of dialysed phage was added, followed by brief mixing of the measuring cell. The Signal was monitored for 2 h and extracellular K^+^ concentration was calculated using the standard curve. A BW25113 *ΔlamB* mutant strain that lacks the receptor for λ adsorption was used as a negative control, and the experiment was performed in biological triplicates.

### Purification of recombinant proteins

N-terminally hexa-His-tagged BREX or Ocr proteins were expressed in BW25113 cells transformed with pBAD-His_6_-BrxN/Ocr or the corresponding BrxX^Y511A^ derivative. Since the presence of N-His in BrxX, and N-His-TEV in Ocr did not interfere with the *in vivo* or *in vitro* functions of the protein, a non-processed variants were used throughout the work. Overnight culture obtained from the freshly transformed cells was diluted 100x into LB media (8 L) supplemented with ampicillin, grown at 37°C until OD_600_ reached 0.9, and induced with L-arabinose (0.2% w/v final concentration). Following overnight expression at 18°C, bacteria were harvested by centrifugation (10 000 g, 4°C), and the pellet was resuspended in a buffer A (50 mM Tris-HCl pH 8.0, 300 mM NaCl, 20 mM imidazole, 5 mM 2-mercaptoethanol, 5% glycerol) supplemented with a cOmplete Protease Inhibitor Cocktail (Roche). Cells were lysed by sonication and the lysate was centrifuged (21 000 g, 4°C). The clarified lysate was applied to 5 ml HisTrap HP columns (Cytiva) using an ӒKTA pure FPLC system (Cytiva). The resin-bound protein was first washed with 8 column volumes (CV) of buffer A, followed by gradient elution with buffer B (50 mM Tris-HCl pH 8.0, 500 mM NaCl, 500 mM imidazole, 5 mM 2-mercaptoethanol, 5% glycerol). BrxB and BrxZ were dialysed overnight at 4°C against GF buffer (50 mM Tris-HCl pH 8.0, 150 mM NaCl, 5 mM 2-mercaptoethanol) and concentrated for gel-filtration (see below) directly after this step, whilst BrxA, BrxC, BrxX, BrxX^Y511A^, BrxL and Ocr were first purified on a heparin affinity column. For this, fractions corresponding to the protein peak were pooled and carefully diluted with buffer C (50 mM Tris-HCl pH 8.0, 5 mM 2-mercaptoethanol, 5% glycerol) until the concentration of NaCl reached 100 mM. The sample was loaded onto a 5 ml HiTrap Heparin column (Cytiva) and washed with 5 CV of buffer C supplemented with 100 mM NaCl. Proteins were eluted by a gradient of buffer D (50 mM Tris-HCl pH 8.0, 5 mM 2-mercaptoethanol, 1 M NaCl, 5% glycerol). Ocr protein was collected from the flowthrough fractions, while other proteins were bound to the column. Therefore, Ocr was additionally purified via a second pass through a HisTrap HP column. Protein-containing fractions were collected and concentrated/buffer exchanged to GF buffer using 10-, 30- or 100-kDa Amicon centrifugal filter units (Merk), depending on the protein size. Concentrated proteins were loaded onto HiPrep 26/600 Superdex 200 size exclusion column (Cytiva) connected to an ӒKTA pure FPLC system and the run was carried in a GF buffer. Protein-containing fractions were analysed by SDS-PAGE, pooled, concentrated, distributed into 5 µl aliquots and snap-frozen in liquid nitrogen for storage at −80°C for further *in vitro* studies. The protein concentration was measured using 280 nm absorbance with NanoDrop 8000 (Thermo Scientific).

### Electrophoretic mobility shift assay (EMSA)

Oligos used in EMSA are listed in **Supplementary Table S2**. To anneal dsDNA duplexes, equal amounts of 100 μM 5’-Cy5 labelled forward strand and unlabelled reverse strand oligonucleotides (Evrogen/Merck) were mixed in the annealing buffer (10 mM Tris-HCl pH 8.0, 50 mM NaCl). After heating to 95°C for 10 min, the mixture was allowed to slowly cool to room temperature, before dilution in mQ to achieve 100 nM final stock concentration. *BREX proteins binding to non-specific and specific DNA substrates.* Proteins (in a range of concentrations), supplemented with 0.5 mM SAM or SAH, if necessary, were mixed with 20 nM Cy5 labelled DNA in EMSA buffer (0.5x TBE pH 8.0, 35 mM NaCl, 5% glycerol) in final 12 μl reactions and incubated for 30 min at room temperature. A substrate with methylated BREX site was ordered through Sigma.

*EMSA of BrxX and DNA in the presence of Ocr.* The procedure was the same as above, 750 nM BrxX was incubated with either 20 nM DNA (bearing one BREX site) or Ocr for 45 min at room temperature, followed by addition of either Ocr or 20 nM dsDNA and incubation for further 45 min at room temperature. Ocr molar concentration was calculated for the Ocr dimer.

*Competitive binding of Ocr and DNA to BrxX.* 400 nM BrxX was incubated with 20 nM 43bp Cy5-dsDNA (bearing one BREX site) for 20 min at room temperature, followed by addition of either Ocr or unlabelled 43bp dsDNA (bearing one BREX site) and incubation for 20 min at room temperature. Ocr molar concentration was calculated for the dimer.

All binding reactions were run on a 10% polyacrylamide (37.5:1) TBE gels for 35 min at 110V at room temperature. Cy5 labelled DNA was visualized with the Molecular Imager Gel Doc XR System (BioRad).

### Strep-Seq analysis of BrxX binding to T7 DNA

To detect *in vivo* BrxX methyltransferase binding to the phage DNA in a course of infection, a variant of the ChIP-Seq procedure was developed. Cells carrying pBREX AL BrxX C-Strep were grown in 100 ml LB at 37°C till OD_600_ = 0.5. 40 mL aliquot was taken and mixed with the phage T7 ^43^ at MOI=1, infection proceeded for 15 min. Formaldehyde was added to the culture at a final concentration of 1% and cross-linking was carried for 5 min at room temperature. Formaldehyde was quenched by adding glycine to a final 500 mM concentration and the mixture was agitated at 4°C for 20 min. Cells were collected by centrifugation (6 000 g, 10 min, 4°C) and kept at −20°C overnight. Cells were resuspended in lysis solution (StrepA buffer supplemented with 300 μg/ml lysozyme) and incubated at 37°C for 30 min following sonication disruption (65% power, 10 sec pulse, 20 sec pause, 12 cycles on Qsonica sonicator with 2 mm sonotrode). Sonication conditions were optimised to produce DNA fragments of 200-600 bp mean shear size. Samples were treated with RNAse A and 1/20 volume was taken for the purification of total DNA which served as mock DNA control without enrichment. The rest of the sample was loaded on the StrepTrap HP chromatography column to purify Strep-tagged BrxX. The eluted fraction was kept at 65°C to reverse formaldehyde cross-links. DNA was purified with phenol:chloroform method and sequenced on MiniSeq platform (Illumina) with paired-end 150 cycles (75 + 75). Reads from StrepTrap enriched sample and mock DNA sample were mapped to the T7_fusion_ genome using bwa mem. Genome coverage was calculated and analysed with bedtools, genomecov, and samtools and was further normalized to the sequencing depth and the genome size to achieve CPM values. Coverage was visualized using seaborn package in Python as the ratio of the enrichment to the mock sample CPM values.

### Biolayer Interferometry (BLI)

Oligos used in BLI are listed in **Supplementary Table S2**. To anneal dsDNA duplexes, equal amounts of 100 μM 5’-biotinylated forward strand and unlabeled reverse strand oligonucleotides (Syntol) were mixed in the annealing buffer (10 mM Tris-HCl pH 8.0, 50 mM NaCl). After heating to 95°C for 10 min, the mixture was allowed to slowly cool to room temperature. All kinetic assays with Octet R2 were set up in 96-well plate format using 200 μL reaction volumes and were performed in running buffer containing 20 mM Tris-HCl pH 8.0, 100 mM NaCl, 0.05 mg/mL BSA, and 0.02% Tween-20 at 30 °C with the orbital shake speed of 1000 rpm. Reactions were supplemented with 0.5 mM SAM in appropriate measurement series.

Kinetic assays were performed by first capturing 225 nM of 5’ biotinylated dsDNA (20 or 40 bp DNA containing either the BREX site or polyC sequence) onto SAX Octet biosensor in running buffer. The initial baseline was generated for 60 sec, loading of the bait was performed for 100 sec, and a second baseline was generated for 150 sec. Association with BrxX analyte (0.05 – 1.6 μM) was performed for 150 sec and final dissociation was performed for 450 sec. For Ocr competitive binding assay, the initial baseline was generated for 30 sec, loading of 40 bp DNA with one BREX site was performed for 100 sec, a second baseline was generated for further 150 sec, association was performed for 7 min in wells filled with 1.5 μM BrxX analyte supplemented with either untagged 40 bp dsDNA or Ocr, followed by further 21 min dissociation in running buffer. Ocr molar concentration was calculated for the dimer. In each run, one of the SAX sensors loaded with biotinylated dsDNA was incubated in a running buffer without the analyte (BrxX) to capture the reference signal. It was also validated that incubation of the SAX sensor with BrxX doe does not result in a binding signal. Fresh SAX biosensors were used without any regeneration step.

The raw data measured on Octet R2 was analysed using Octet Analysis Studio software (version 12.2.0.20). After single reference subtraction, binding sensorgrams were first aligned at the beginning of the association cycle, inter-step corrected to the association step, and filtered according to the Savitzky-Golay algorithm. Binding sensorgrams were globally fit to a 1:1 Langmuir binding model for 20 bp substrates or to 2:1 heterogenous binding model for 40 bp substrates. Some curves with low protein concentrations were excluded from the analysis due to the lack of response signal.

### Cryogenic electron microscopy (cryoEM) sample preparation

BrxX-DNA complex was prepared containing 250 mM DNA (43 bp, H2T), 50 mM BrxX monomer and 1 mM SAM. The mixture was dialyzed overnight at 4 C into cryoEM buffer (20 mM HEPES-NaOH pH 7.5, 2.5 mM MgOAc, 150 mM NaCl and 0.5 mM TCEP) containing 50 µM SAM. For BrxX-Ocr, 10 mg/ml of reconstituted BrxX-Ocr complex was dialyzed overnight at 4 °C into cryoEM buffer. Following dialysis, SAM was added to 1 mM and CHAPSO to 8 mM final concentrations. Both complexes were spun for 1 hour at 15 900 g prior to grid preparation to remove any potential aggregates. Aliquots of 3.5 µl of BrxX:DNA or BrxX/Ocr complexes were applied to glow-discharged (Leica, 60 s/8 mA) Quantifoil holey carbon grids (R2/1, 300 copper mesh). After 30 s of incubation with 100% chamber humidity at 4° C, the grids were blotted for 3.5 s (BrxX-Ocr) or 4.5 s (BrxX-DNA) and plunge-frozen in liquid ethane using a Vitrobot mark IV (FEI).

### Single-particle cryoEM data collection

*Initial screening & analysis.* For initial screening, data were collected on Talos microscope (Thermo) equipped with a Gatan Elsa side-entry holder & Falcon 4i direct electron detector operated at 200 kV at JIC BioImaging facility. Datasets of 4000-5000 movies in EER mode were collected and analysed before progressing to Krios microscope data collection. *BrxX:DNA.* CryoEM data were collected at Astbury Biostructure Laboratory (ABSL) Cryo-EM facility (University of Leeds, UK) on Krios G2 microscope (Thermo) operated at 300 kV and nominal magnification of 165 kx. Movies were recorded in counting mode on a Falcon 4i direct electron detector (Thermo) in EER format using EPU v 3.6. Movie frames were collected at the calibrated physical pixel size of 0.74 Å/px with a defocus range set to −3 µm to −0.9 µm. A dose rate of and exposure time was chosen to result in a total dose of ∼50 electrons/Å^2^. Full statistics for cryoEM data collection are listed in **Supplementary Table S3**.

*BrxX:Ocr.* CryoEM data were collected at Astbury Biostructure Laboratory (ABSL) Cryo-EM facility (University of Leeds, UK) on Krios G2 microscope (Thermo) operated at 300 kV and nominal magnification of 120 kx. Movies were recorded in counting mode on a Falcon 4i direct electron detector (Thermo) in EER format using EPU v 3.3. Movie frames were collected at the calibrated physical pixel size of 0.68 Å/px with a defocus range set to −3 µm to −0.9 µm. A dose rate of and exposure time was chosen to result in a total dose of ∼35 electrons/Å^2^. Full statistics for cryoEM data collection are listed in **Supplementary Table S3**.

### CryoEM data analysis

*BrxX:DNA:* All processing was done in cryoSPARC^69^ v.4.4+. 11 152 movies were motion and CTF corrected in patch mode. After manual curation, 9 980 movies were retained for further analysis. Particles were picked with Topaz^70^ (trained on a small set of manually-picked micrographs) and 2×2 binned particles (610 100) were subjected to a single round of 2D classification (100 classes, 40 iterations, 20 full iterations, batchsize 300, max resolution 5). 265 295 retained particles displaying high-resolution features underwent 3D classification with 2 classes (*ab initio*, 219 641 particles retained) and was refined to 3.03 Å resolution. Particles were re-extracted at the original pixel size using updated coordinates. Non-uniform refinement^71^ of this particle set with local CTF correction resulted in a 2.47 Å map. To further improve the resolution, particles underwent local reference-based motion correction (‘Bayesian polishing’) as implemented in cryoSPARC v.4.4+^72^. Two rounds of particle polishing were carried out followed by non-uniform refinement with local defocus refinement, global CTF refinement (tilt, trefoil, spherical aberration and anisotropic magnification^73^) and Ewald sphere correction^74^; this resulted in a final map with 2.2 Å global resolution as measured by cryoSPARC.

*BrxX:Ocr:* Processing was done in cryoSPARC v 4.1+. 5 100 collected movies were motion & CTF-corrected and manually curated. 4 678 movies were retained for further analysis. 24 micrographs were picked manually and used to train Topaz. Topaz-picked particles (244 977) were used to create an initial map of BrxX_2_:Ocr_2_ complex. This map was used to create equally-spaced 2D projections to use as templates, and particles were more thoroughly picked using cryosparc template picker (396 923) and extracted as twice binned (1.36 Å/pix). Based on manual analysis of 2D class averages, 227 907 particles were selected that represented a BrxX_2_:Ocr_2_ complex with a minor set of particles assigned to BrxX apo (38 496, see below). For 227 907 particles (main set), *ab initio* 3D classification was used to further separate low-resolution particles. 211 818 particles representing complete complex were retained, and by means of heterogenous refinement (batch size 9990, final iterations 20) separated into two classes of almost equal size (114 826 and 96 992 particles) that differed by the slight change in the orientation of the ‘upper jaw’ of the better-resolved BrxX monomer (see **Supplementary Figure S5**). A larger class was retained for more detailed analysis. After a NU-refinement, these particles were classified without alignment using a mask around the ‘upper jaw’ (MTD+NTD). Parameters for 3D classification job were: 5 classes, 10000 batch size, 20 O-EM epochs, target resolution 4 Å. Four best classes were retained (94 764 particles). Particles underwent one round of Bayesian polishing as implemented in cryoSPARC, followed by local CTF refinement, global CTF refinement (tilt, trefoil, spherical aberration and anisotropic magnification) and Ewald sphere correction. Final consensus refinement resulted in a 2.91 Å map with the resolution being highest for the Ocr dimer and two BrxX TRDs. To improve the resolution around the ‘upper jaw’ (MTD+NTD), local refinement was carried out using the corresponding mask, resulting in the 2.84 Å focussed map for the MTD+NTD of a single BrxX monomer within the complex. For the particles representing BrxX apo, *ab initio* 3D classification was carried out with 2 classes. The best class (24 284) was refined to 3.88 Å.

### Model building and refinement

*BrxX:DNA:* AlphaFold2-generated model for BrxX was used as a starting point. The model was split into four separate domains that were first manually rigid-body fitted in ChimeraX^52^, followed by manual re-building in Coot^75^ as necessary. DNA initial model was built using cryoREAD server^76^, followed by manual adjustment in Coot using local restraints. Real-space refinement was performed in phenix.refine^77^ (using Ramachandran restrains, and secondary structure restraints for protein and DNA). *BrxX:Ocr*: A refined BrxX:DNA model was split in two halves corresponding to the ‘upper’ and ‘lower’ jaw (1-660 and 661-1205) which were rigid-body fit into the density for the best resolved monomer. An Ocr-inhibited *E. coli* RNAP structure (PDB: 6R9G) was used to generate an Ocr initial model which was rigid-body fit into the density. Missing elements were manually built in Coot; Ramachandran restraints were used to built loops with weaker density, particularly residues 422-430. After building best resolved BrxX monomer, it was copied and rigid-body fitted into the density for the second monomer, using a well-resolved two-helix bundle at the tip of the CTD for guidance. NCS restraints were used in Phenix.refine to stabilise first rounds of refinement, and were switched off during later rounds as the structure is not fully symmetrical. Ramachandran restrains, and secondary structure restraints were used. *BrxX apo*: a well-resolved BrxX monomer from the BrxX:Ocr structure was rigid body fit in the map, and refined in phenix.refine using Ramachandran and secondary structure restraints. MolProbity^78^ and MTriage^79^ were used to validate all models and maps. Statistics for the final models are reported in **Supplementary Table S3**.

### BREX methylation restriction-sensitivity assay

Plasmid pHERD30t with and without overlapping AluI/BREX site was purified from BREX^+^ (*E. coli* HS) or BREX^-^ cells (*E. coli* BW25113) using Monarch Plasmid Purification Kit (NEB). 100 ng of plasmid was restricted with AluI (ThermoScientific) according to manufacturer’s instructions. Non-cleaved and cleaved plasmids were run at 1% agarose gel in 1x TAE buffer and stained with ethidium bromide.

### *In vitro* methylation assay

BrxX or Dam protein (100 μM final concentration) was mixed with 5 µg of the annealed 40 bp DNA substrate, containing overlapping BREX/AluI site and a single Dam site, in methylation buffer (20 mM Tris-HCl pH 7.5, 100 mM NaCl, 0.1 mg/ml BSA, 1 mM DTT) optionally supplemented with 2 mM SAM. Reaction was incubated overnight at room temperature. DNA was purified with phenol-chloroform, skipping the 70% ethanol washing step^80^. 350 ng of purified DNA was restricted (according to manufacturer’s instructions) with DpnI (NEB) or DpnII (NEB) to monitor Dam site methylation status or with AluI (ThermoFisher) to monitor BREX site methylation status. The resulting fragments were run on 12% polyacrylamide gels (37.5:1) in 1x TAE buffer. The gels were visualized with SYBR Gold (ThermoFisher).

### HPLC-MS analysis of nucleosides

1 µg of purified dsDNA substrate incubated with Dam or BrxX, as described above, was digested with Nucleoside Digestion Mix (NEB) at 37°C overnight. Nucleosides were loaded onto Agilent Poroshell 120 SB-C18 column (4.6 x 100 mm, 2.7 µm) and were analyzed on Agilent 1200 HPLC-MS system with ESI source and Q-TOF detector (Agilent). Gradient conditions were as following: solution A - 5 mM ammonium acetate, pH = 5.3; solution B - 90% acetonitrile; LC run was carried at 40°C, 0.3 ml/min speed and 1 µl of sample was loaded. The column was washed for 5 min with 2% B, followed by linear increase to 30% B till 30 min, linear increase to 100% B till 36 min, and linear decrease to 2% B till 40 min. UV detection was carried at 260 nM. LC-MS/MS data were analysed in MassHunter, nucleosides and their modified variants were searched in EICs of expected m/z values for nucleosides and dm6A.

### Efficiency of plating (EOP) assay

To determine the titer of active phage particles in cell lysates, the double agar overlay method was used. Overnight culture of bacteria (100 μl) was mixed with 10 ml of 0.6% top LB agar supplemented with 0.2% maltose, 5 mM MgSO_4_ and appropriate antibiotics and poured on the surface of 1.2% LB agar plates. 10 μl drops of serial 10-fold phage lysate dilutions were spotted on the top agar, allowed to dry and plates were incubated at 37 °C overnight. The level of protection was determined as the ratio of phage titers obtained on a non-restrictive (BREX^−^) host relative to that on restrictive (BREX^+^) host. All experiments were performed in biological triplicates.

### *In vivo* estimation of BREX methylation via phage λ sensitivity assay

Estimation of the *in vivo* efficiency of BREX methylation was conducted via EOP assay with the phage λ induced from BREX^-^, BREX^+^ or other indicated culture. The results of this assay correlate with direct estimation of BREX methylation with PacBio sequencing, as shown in our previous works^18,43,46^. Estimation of the pBREX AL BrxX^Y511A^ methylation activity was performed with *λ_ts_*, as described before^18,43,46^. However, *λ_ts_* phage was not compatible with the plasmids used for expression of BREX proteins combinations due to the presence of the *Cm^R^* gene. Therefore, for the determination of the minimal set of proteins required for BREX methylation, we used BW25113 *λ_w_*_t_ lysogen. *λ_w_*_t_ lysogen was obtained by infecting BW25113 with *λ_w_*_t_ in liquid culture and selecting colonies resistant to *λ_w_*_t_ superinfection. Lysogenic cells were transformed with pBTB-2 (BREX^-^ culture), pBREX AL (BREX^+^ culture), pBAD BrxX or a set of plasmids expressing different combinations of BREX proteins^19^: BCXZ (pBAD BrxX + pBREX1 ΔBrxA + pTG BrxZ), CXZ (pBREX2 ΔBrxL + pTG BrxС) or BrxX+N (pBAD X + pTG N, where N is any other BREX protein). Overnight cultures were diluted 100x in 35 ml LB with 0.2% arabinose to induce BREX proteins expression and appropriate antibiotics and grown at 37°C until OD_600_ ∼ 0.4. To induce *λ_w_*_t_ lysogen cells were treated with UV. First, to increase transparency of the culture, cells were harvested by centrifugation (6 000 g, 20°C, 10 min) and washed twice in an equal volume of SM3 buffer (see above) and then in 10 ml SM3 buffer. Suspension of cells was decanted into sterile 100×15mm Petri dish, which was placed without the lid on a shaking platform under the UV lamp (30W) at a distance of ∼50 cm. Suspension was exposed to UV for 30 sec. After that, cells were collected in a tube and harvested by centrifugation (6 000 g, 20°C, 10 min). Cell pellet was resuspended in 5 ml LB with 0.2% arabinose and incubated at 37°C for 3 h until the visible lysis. Cell debris was removed by centrifugation and lysate was cleared with chloroform, as described elsewhere^81^. The methylation status of collected *λ_w_*_t_ was measured by determining the ratio of phage titer obtained on a non-restrictive (BREX–) host relative to that on restrictive (BREX+) host.

### Toxicity assay

BREX^-^, BREX^+^ or pBREX AL BrxX^Y511A^ potential toxicity was estimated in a spot-test assay. Cells were grown in 10 ml of LB and diluted to OD_600_=0.6, followed by serial 10-fold dilution droplets plating on 1.2% LB-agar supplemented with required antibiotics.

### *In vivo* protein pull-down and size exclusion chromatography

Overnight cultures of *E. coli* strains with plasmids encoding Strep-tagged BREX proteins in a context of the full BREX cluster^43^ (pBREX AL BrxN C-Strep, where N is any of the BREX proteins) were diluted 100-fold in 3 L of LB media and incubated overnight at 18°C in Thomson Scientific Ultra Yield flasks with aerated lids (1.5 L per flask). Cells were harvested by centrifugation at 4 000 g at 4 °C for 30 min after reaching OD_600_ ∼ 0.9. Pellets were washed in 50 ml of StrepA buffer (150 mM NaCl, 1 mM EDTA, 5 mM 2-mercaptoethanol, 100 mM Tris–HCl pH 8.0), and resuspended in the same buffer supplemented with a cOmplete Protease Inhibitor Cocktail (Roche) and lysozyme (0.2 mg/mL). Cells were disrupted by sonication on ice, and the lysate was clarified by centrifugation at 21 000 g, 4 °C for 30 min. Strep-tagged proteins were purified on two stacked 5 mL StrepTrap HP (Cytiva) columns, connected to the NGC Chromatography System (BioRad). Protein-containing fractions were concentrated using 3 kDa Amicon centrifugal filter units (Merk) and analyzed by SDS-PAGE. The identity of protein bands was determined by matrix-assisted laser desorption/ionization time-of-flight (MALDI-TOF) mass spectrometry. Samples were prepared with Trypsin Gold (Promega) in accordance with manufacturer’s instructions. Mass spectra were obtained using the rapifleX system (Bruker). The molecular weight of protein complexes was determined by size exclusion chromatography performed on Superdex 200 Increase 10/300 column (GE Healthcare) calibrated with High Molecular Weight calibration kit (GE Healthcare). Protein fractions were checked for DNA presence by incubation with DNAse I at 37°C and electrophoresis in 1% agarose 1xTAE gel and ethidium bromide staining.

### Protein stability estimation with a label-free thermal shift analysis

To estimate BrxX and BrxX^Y511A^ thermostability, 7 µM of protein was mixed with 0.3 mM either SAM or SAH or was incubated without co-factors at room temperature for 10 min in a 10 µl buffer (20 mM Tris-HCl pH 8.0, 100 mM NaCl). Mixture was loaded into Tycho NT.6 capillaries and protein stability was monitored by fluorescence (as a ratio of intrinsic tryptophan to tyrosine fluorescence detected at 350 nm and 330 nm, correspondingly) upon a gradient temperature increase (as a 30°C/minute temperature ramp applied from 35 to 95°C).

### Quantification and statistical analysis

All toxicity and EOP assays were performed at least in three biological replicates. All values are presented in the form of a mean +/-standard deviation. EMSAs and *in vitro* reactions with AluI-containing dsDNA substrates were performed in duplicates and representative EMSA gels and HPLC profiles are presented. All calculations were performed in MS Excel.

---

**Supplementary Figure S1.**
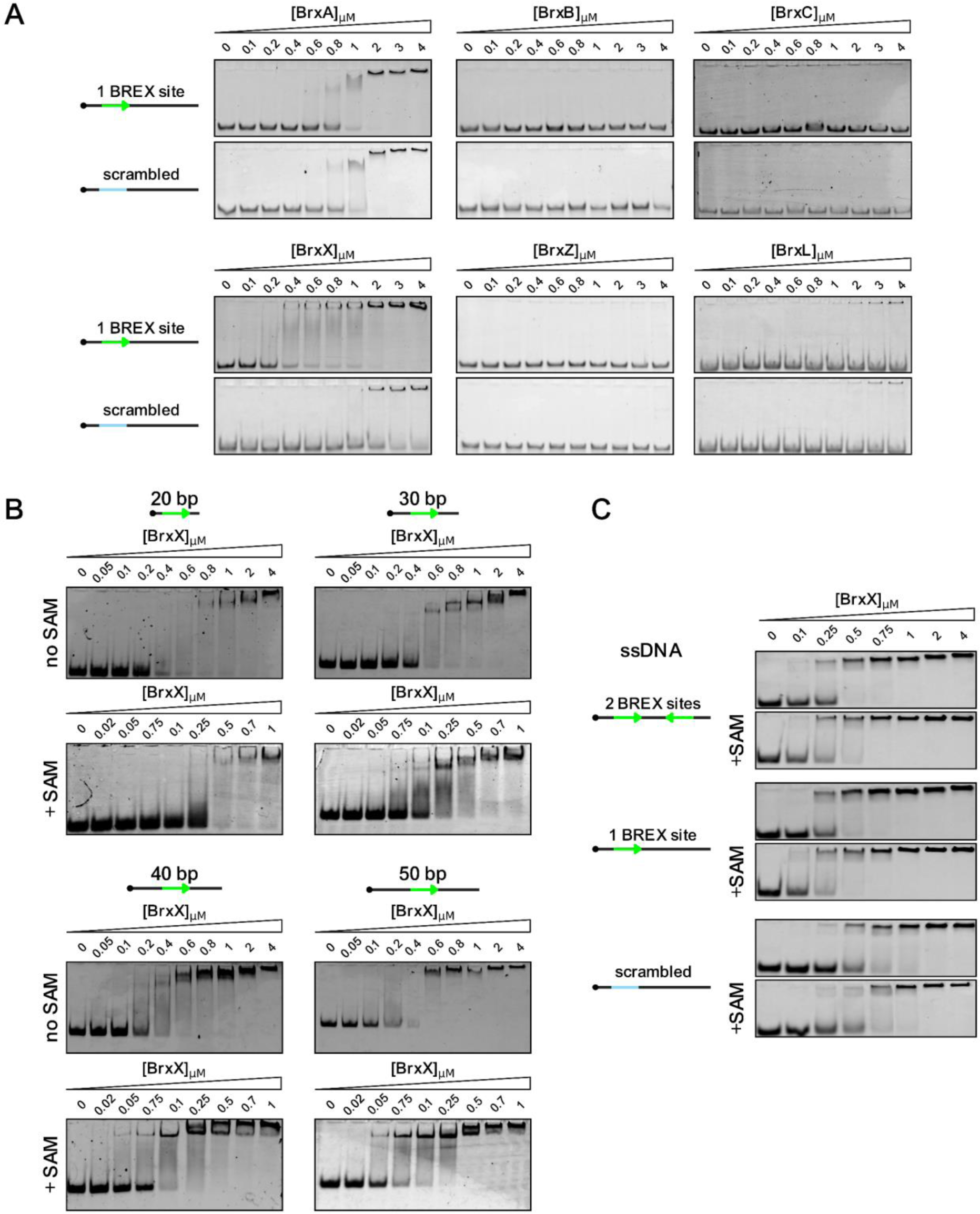
BREX proteins interact with DNA substrates. (**A**) EMSA with 20 nM Cy5-labelled 43 bp dsDNA substrate without BREX sites (scrambled) or with one BREX site, incubated with indicated amount of BREX proteins. (**B**) EMSA with 20 nM Cy5-labelled dsDNA substrates of different length bearing one BREX site, incubated with indicated amount of BrxX without co-factors or in the presence of 0.5 mM SAM. (**C**) EMSA with 20 nM Cy5-labelled 43 bp ssDNA substrate without BREX sites (scrambled), with one, or two BREX sites, incubated with indicated amount of BrxX without co-factors or in the presence of 0.5 mM SAM. Representative gels from triplicate experiments are shown.

**Supplementary Figure S2.**
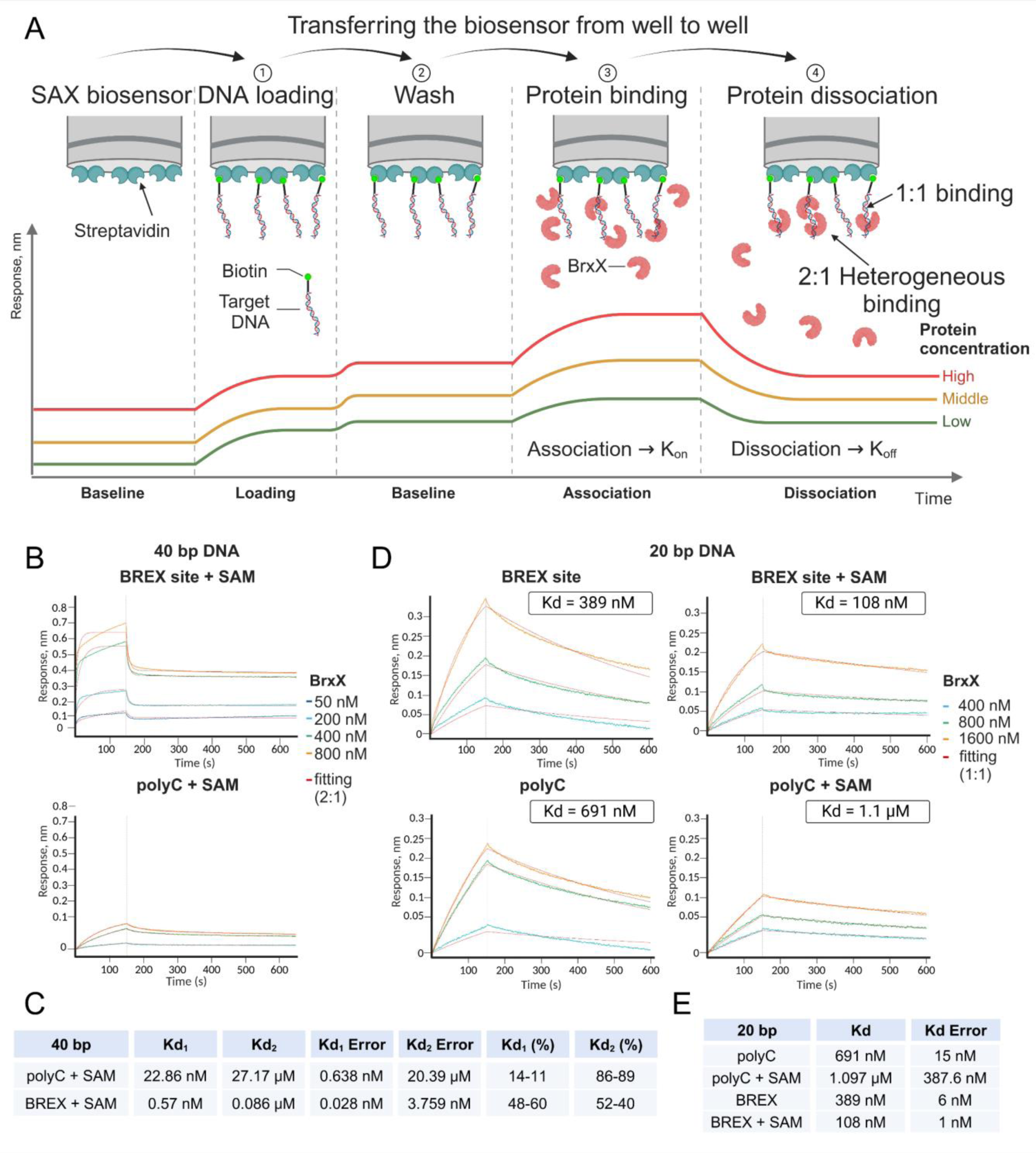
Biolayer Interferometry (BLI) measurement of BrxX:DNA interaction. (**A**) A principle of the BLI assay, measuring BrxX interaction with biotinylated dsDNA substrate. Examples of kinetic curves are provided. Two binding modes (1:1 and 2:1) describing possible BrxX:DNA interactions are presented. (**B**) Association and dissociation kinetics of BrxX interacting with a 40 bp biotinylated dsDNA substate bearing one BREX site or a polyC sequence. 0.5 mM SAM was present when indicated. Determined kinetic parameters are shown in (**C**) and the data was fit to the 2:1 binding model. (**D**) Association and dissociation kinetics of BrxX interacting with a 20 bp biotinylated dsDNA substate bearing one BREX site or a polyC sequence. 0.5 mM SAM was present when indicated. Determined kinetic parameters are shown in (**E**) and the data was fit to the 1:1 binding model.

**Supplementary Figure S3.**
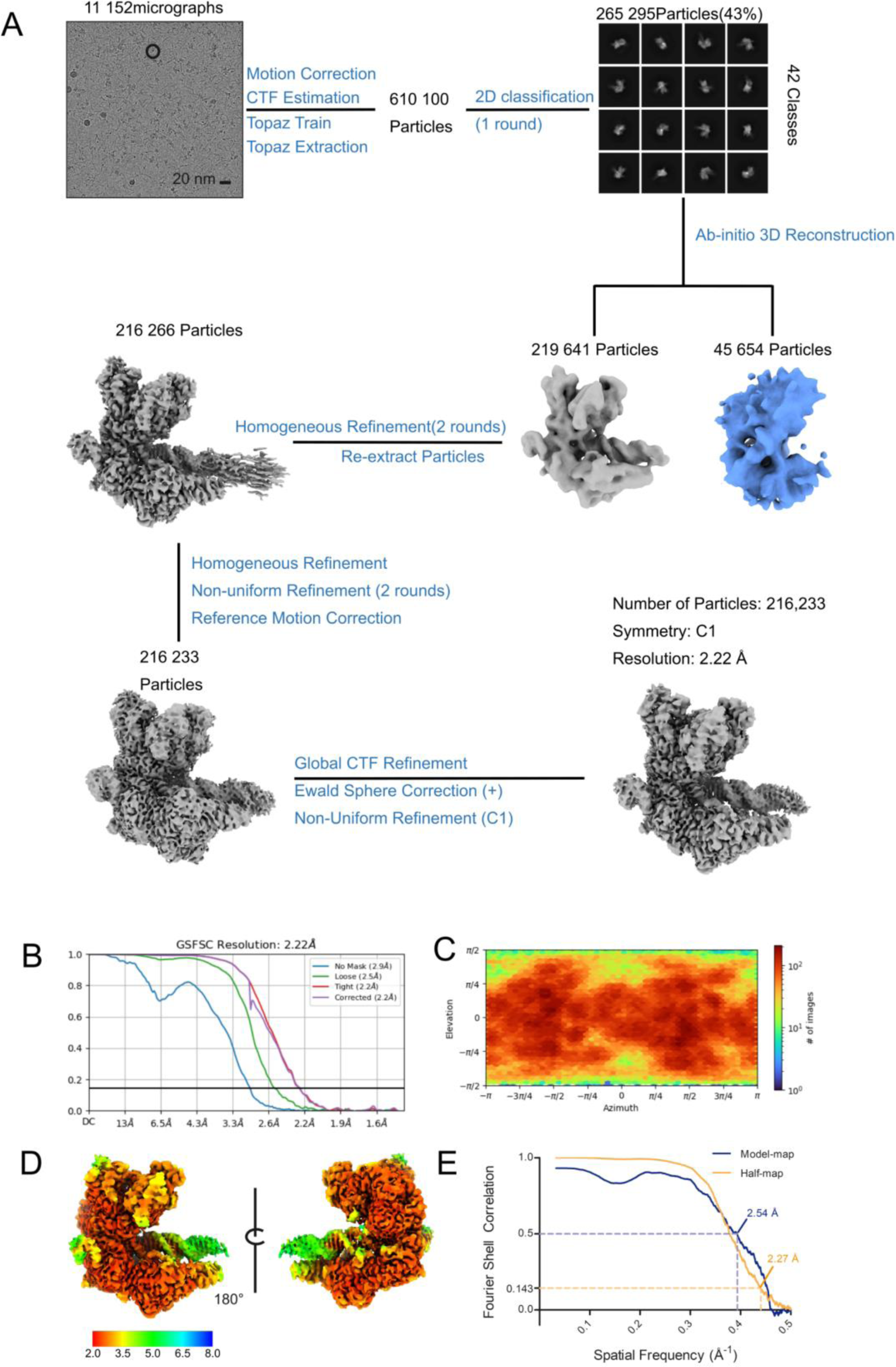
CryoEM data processing for BrxX:DNA. (**A**) A representative motion-corrected micrograph and the processing scheme (see *Methods* for description). (**B**) FSC curve for the final reconstruction as output by cryoSPARC. (**C**) Euler angle distribution as output by cryoSPARC. (**D**) Local resolution maps illustrating resolution distribution. (**E**) Map-to-model FSC curve (blue) compared with half-maps FSC curve (orange) as output by Phenix.mtriage (soft mask based on the atomic model is used). FSC=0.5 (for map-to-model FSC) and FSC=0.143 (for half-map FSC) values are indicated with arrows.

**Supplementary Figure S4.**
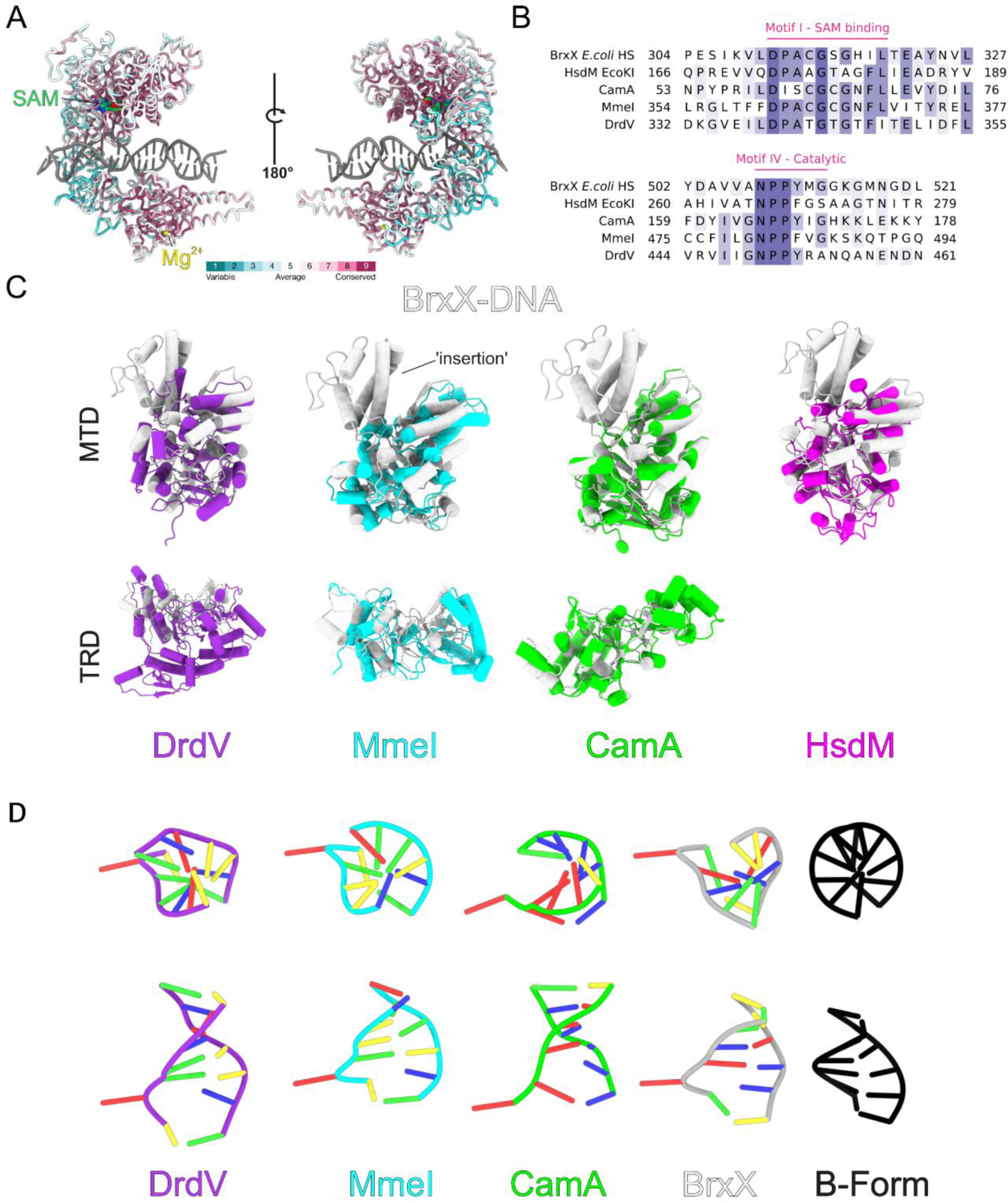
Conservation of different domains of BrxX and comparison with homologous MTases. (**A**) A cartoon representation of a BrxX:DNA model colored by ConSurf^1^conservation scores. SAM molecule and a metal ion (annotated as Mg^2+^) are shown as van der Waals spheres. Note the high degree of conservation around the SAM-binding pocket and the metal presumably associated with recognition of other BREX proteins; TRD has a relatively low level of conservation as a fastest-evolving region. (**B**) Alignment of SAM-binding (Motif I) and methyltransferase catalytic (Motif IV) motifs of BrxX from *E. coli* HS (P0DUF9), Type I R-M EcoKI methylase HsdM (P08957), orphan methylase CamA (Q183J3), and Type IIL R-M methylase-nuclease MmeI (B2MU09). (**C**) Structural alignments of isolated BrxX MTD (*top*) and TRD (*bottom*) domains (white) with homologous domains from restriction-modification systems DrdV (purple, PDB:7LO5), MmeI (cyan, PDB:5HR4) and HsdM (EcoKI; pink; PDB:2AR0) and an orphan Mtase CamA (green, PDB:7LT5). Alignment was done by *matchmaker* algorithm in ChimeraX^2^. (**D**) Comparison of the backbone conformation of DNA between BrxX and the above enzymes. BrxX-bound DNA is most closely resembling DNA in MmeI structure. Ideal B-form DNA is shown in black.

**Supplementary Figure S5.**
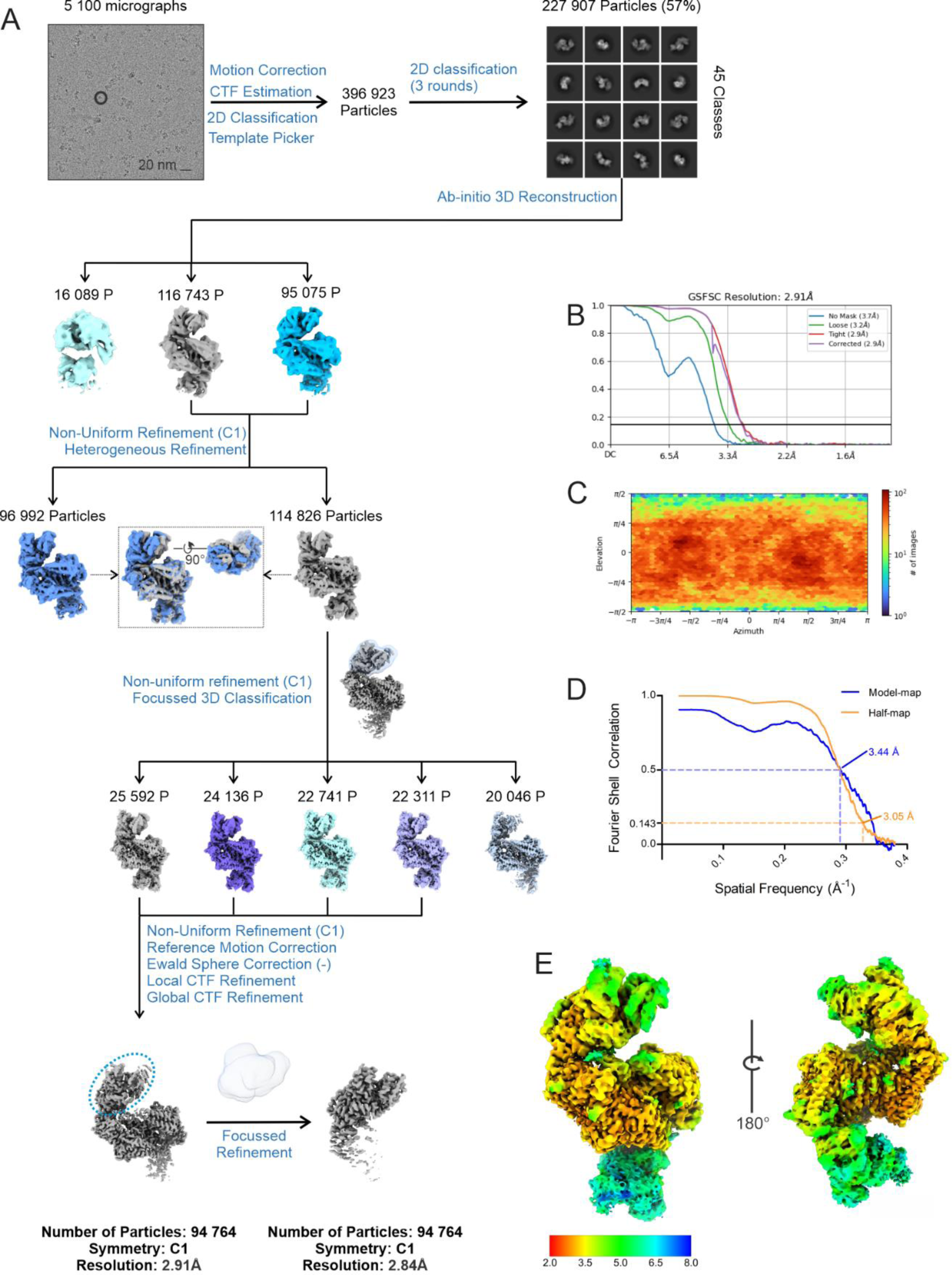
CryoEM data processing for BrxX:Ocr. (**A**) A representative motion-corrected micrograph and the processing scheme (see *Methods* for description). (**B**) FSC curve for the final reconstruction as output by cryoSPARC. (**C**) Euler angle distribution as output by cryoSPARC. (**D**) Map-to-model FSC curve (blue) compared with half-maps FSC curve (orange) as output by Phenix.mtriage (soft mask based on the atomic model is used). FSC=0.5 (for map-to-model FSC) and FSC=0.143 (for half-map FSC) values are indicated with arrows. (**E**) Local resolution maps illustrating resolution distribution.

**Supplementary Figure S6.**
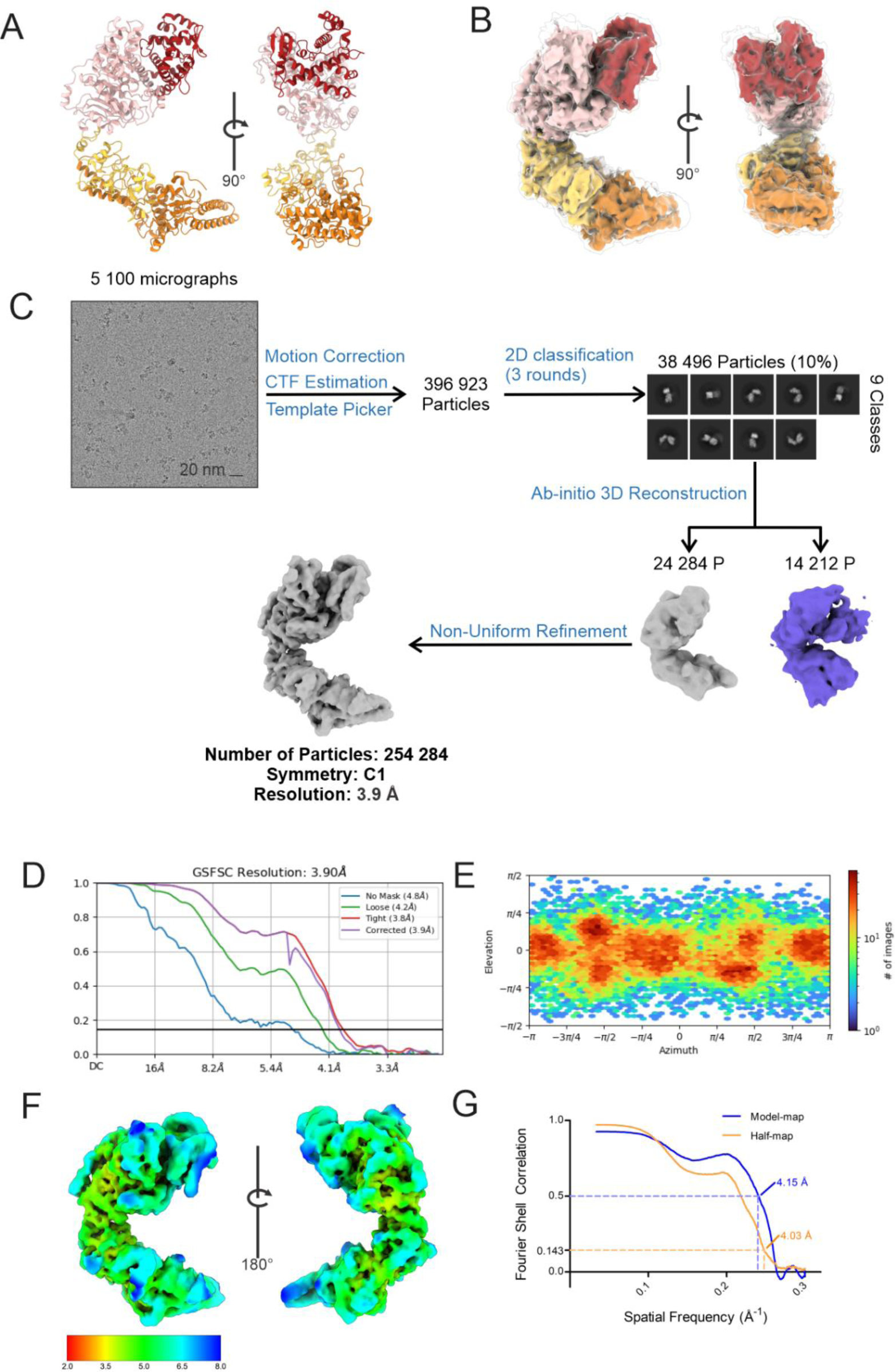
CryoEM data processing for BrxXapo. (**A**) A representative motion-corrected micrograph and the processing scheme (see *Methods* for description). (**B**) FSC curve for the final reconstruction as output by cryoSPARC. (**C**) Euler angle distribution as output by cryoSPARC. (**D**) Map-to-model FSC curve (blue) compared with half-maps FSC curve (orange) as output by Phenix.mtriage (soft mask based on the atomic model is used). FSC=0.5 (for map-to-model FSC) and FSC=0.143 (for half-map FSC) values are indicated with arrows. (**E**) Local resolution maps illustrating resolution distribution.

**Supplementary Figure S7.**
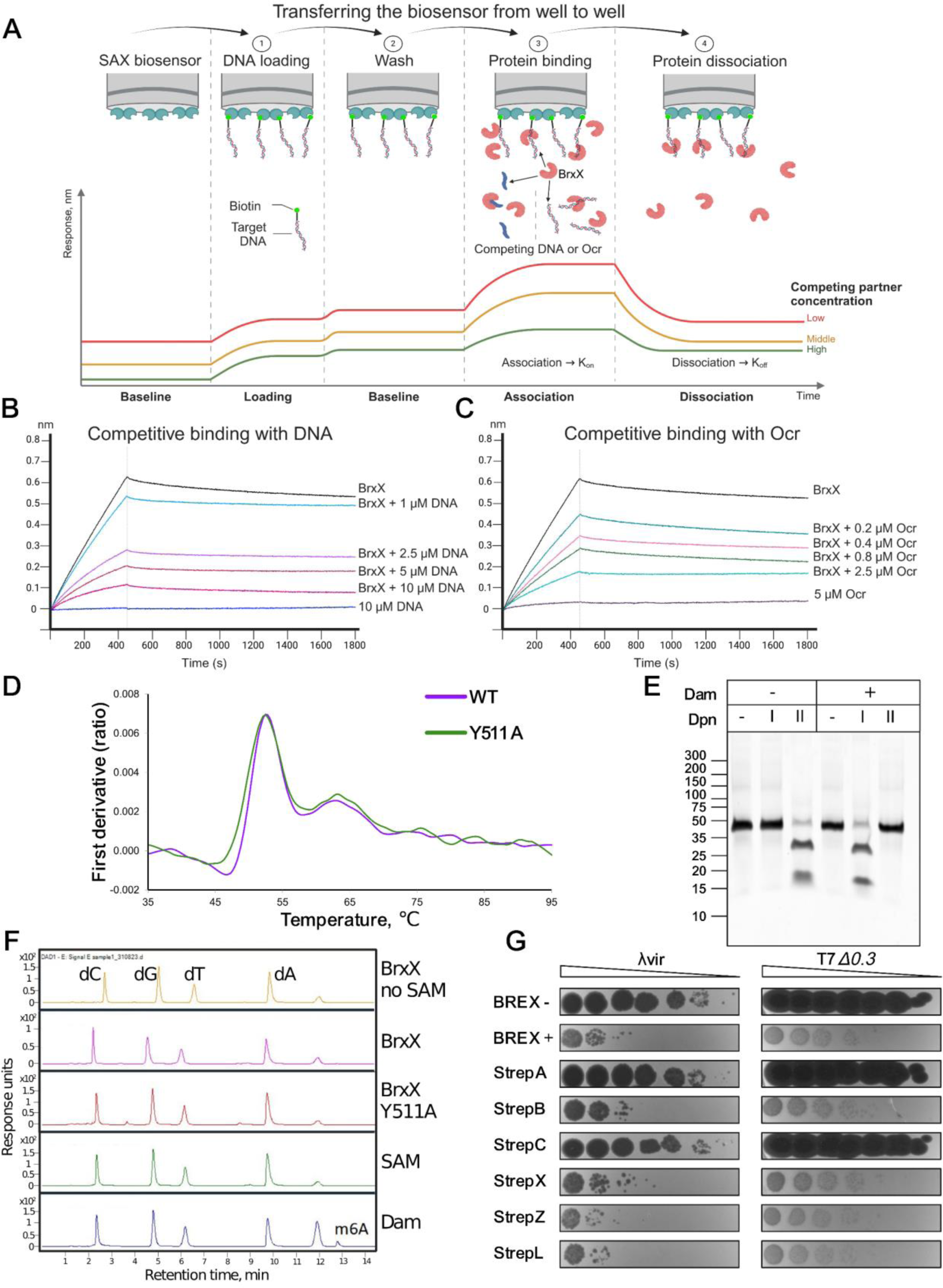
Additional supplementary information for Figures 4 and 5. (**A**) A scheme of the BLI competitive binding assay designed to measure BrxX interaction with biotinylated DNA substrate in the presence of competing non-labeled DNA or Ocr. (**B,C**) Association and dissociation kinetics of 1.5 μM BrxX interacting with a 40 bp biotinylated dsDNA substate bearing one BREX site in the presence of a competing non-labelled dsDNA (**B**) or an Ocr dimer (**C**). (**D**) Thermal stability of BrxX and BrxX^Y511A^ measured with Tycho NT.6. (**E**) DpnI/DpnII restriction-sensitivity assay with 50 bp dsDNA substrate containing a single Dam site and incubated with 100 μM Dam methyltransferase overnight. DpnI cleaves only at methylated Dam sites, while DpnII cleaves only at non-methylated Dam sites. (**F**) HPLC-MS analysis of nucleosides obtained after digestion of a 40 bp dsDNA substrate, bearing an overlapping AluI/BREX site and a single Dam site, incubated with BrxX or Dam methyltransferase overnight. 0.5 mM SAM was present in the methylation reaction, unless otherwise indicated. SAM sample represents a control dsDNA substrate incubated in a SAM-containing buffer without a protein. Peak identity was validated with MS/MS analysis. (**G**) EOP analysis with BREX-sensitive phages *λvir* and T7*Δ0.3* and BREX^-^, BREX^+^ and pBREX AL derivatives bearing Strep-tag (II) coding sequence at the 3’ end of each of the *brx* genes.

**Supplementary Table S1.** Plasmids, bacteriophages and bacterial strains.

| Bacterial strains | Properties | Source |
| --- | --- | --- |
| BW25113 | *E. coli* K12 F<sup>-</sup> $\Delta(araD-araB)567 \Delta lacZ4787(::rrnB-3) \lambda- rph-1 \Delta(rhaD-rhaB)568 hsdR514$ | Lab stock |
| BREX<sup>-</sup> | BW25113 carrying pBTB-2 | 3 |
| BREX<sup>+</sup> | BW25113 carrying pBREXAL | 3 |
| BW25113 $\lambda_{ts}$ lysogen | $\lambda_{ts}$ lysogen of BW25113 | 3 |
| BW25113 $\lambda_{wt}$ lysogen | $\lambda_{wt}$ lysogen of BW25113 | This work |
| QD5003 | Supports lysis by $\lambda cl_{857} S_{am7}$ | Lab stock |
| **Bacteriophages** |  |  |
| T7 | BREX resistant | Lab stock |
| T7$\Delta 0.3$ | *0.3::trxA*, BREX sensitive | 4 |
| T7<sub>fusion</sub> | *0.3* and *0.7* genes fusion in a result of deletion ($\Delta 1256-2735$), leading to the production of semi-functional *Ocr* and increased BREX sensitivity | 4 |
| $\lambda$ | | Lab stock |
| $\lambda_{vir}$ | $\lambda$ mutant with an obligatory lytic lifecycle | Lab stock |
| $\lambda_{ts}$ | *cl<sub>857</sub> bor::Cm* | Lab stock |
| $\lambda_{Sam7}$ | *cl<sub>857</sub> S<sub>am7</sub>*, does not support lysis of the cell | Lab stock |
| **Plasmids** |  |  |
| pBTB-2 | pBBR1 origin, *Kan<sup>R</sup>* | Lab stock |
| pBREX AL | 6 genes *E. coli* HS BREX cluster cloned under native promoter in pBTB-2, *Kan<sup>R</sup>* | 3 |
| pBAD-His<sub>6</sub>-BrxN, N = A, B, C, X, X<sup>Y511A</sup>, Z, L | pBAD-His/B, encoding BrxN protein with N-terminal His<sub>6</sub> tag, *araBAD* inducible promoter, *Amp<sup>R</sup>* | 3 |
| pBAD-His<sub>6</sub>-Ocr | pBAD-His/B, encoding *Ocr* protein with N-terminal His<sub>6</sub> tag and TEV cleavage site, *araBAD* inducible promoter, *Amp<sup>R</sup>* | This work |
| pBREX AL BrxN C-Strep, N=A,B,C, X, Z, L | pBREX AL carrying BrxN with a Strep tag on the C-terminus, *Kan<sup>R</sup>* | 4 |
| pTG N, N = A, B, C | p15A origin, *araBAD* inducible promoter, *Cm<sup>R</sup>* | 3 |
| pBREX 1 $\Delta A$ | pBTB-2 derivative expressing BrxB and BrxC proteins, *araBAD* inducible promoter, *Kan<sup>R</sup>* | 3 |
| pBREX 2 $\Delta L$ | pBAD derivative expressing BrxX and BrxZ proteins, *araBAD* inducible promoter, *Amp<sup>R</sup>* | 3 |
| pBAD BrxX | pBAD-His/B, encoding BrxX protein, *araBAD* inducible promoter, *Amp<sup>R</sup>* | 3 |
| pHERD30t | *E. coli* - *P. aeruginosa* shuttle vector, *Gnt<sup>R</sup>* | Gift of Prof. E. Westra, University of Exeter |
| pHERD30t_BREX/AluI | pHERD30t with an insert of overlapping BREX/AluI sites, *Gnt<sup>R</sup>* | This work |

**Supplementary Table S2.**
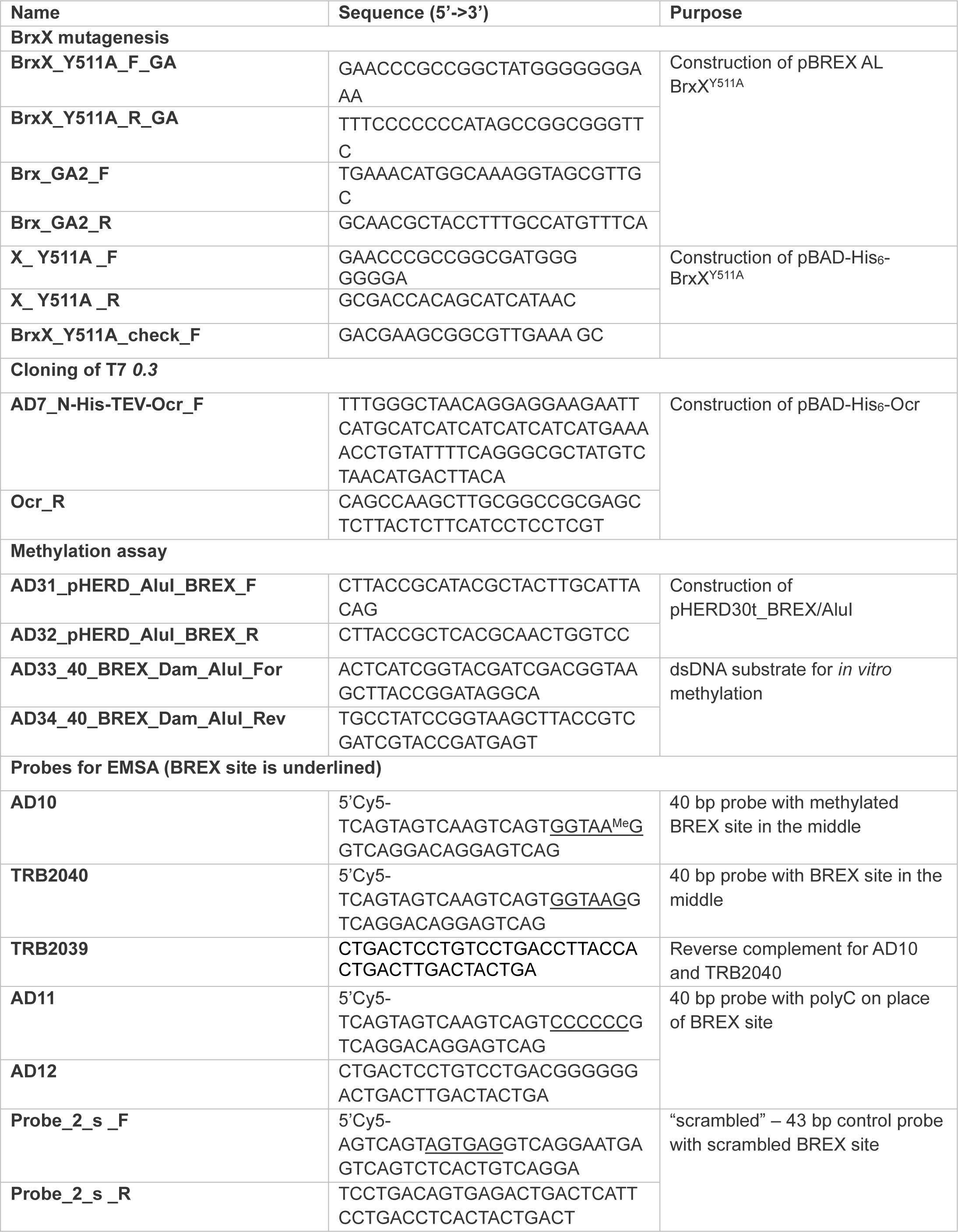
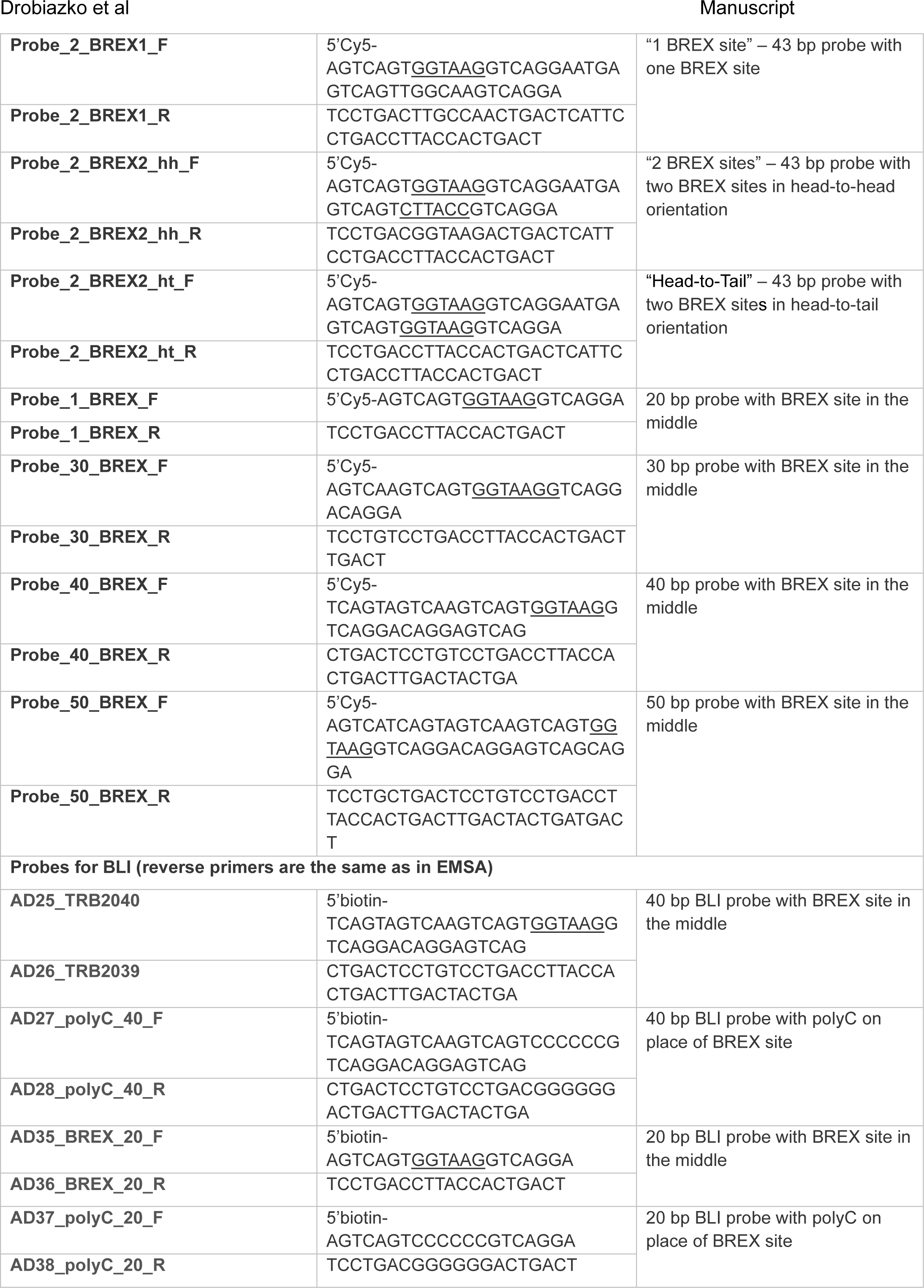
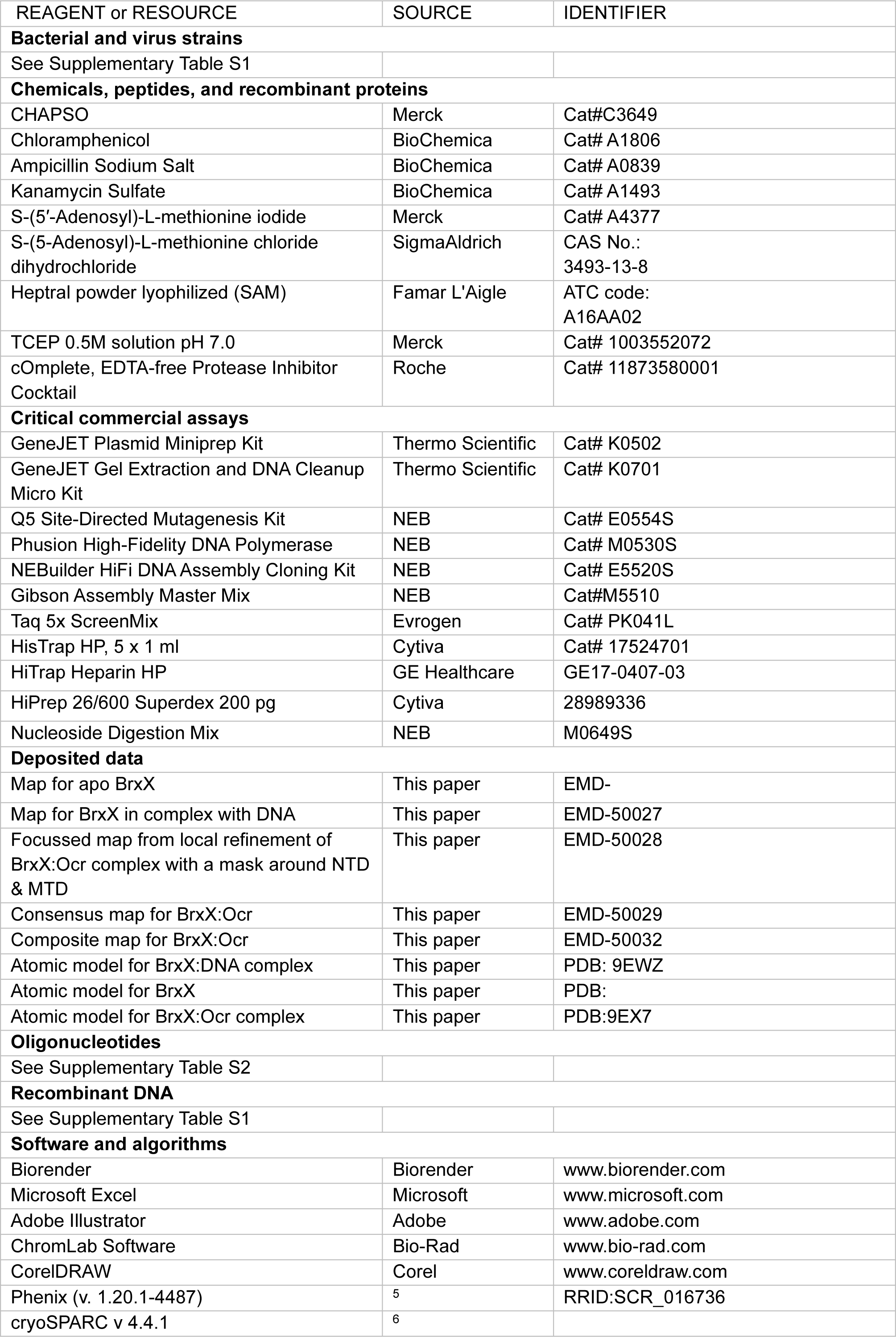
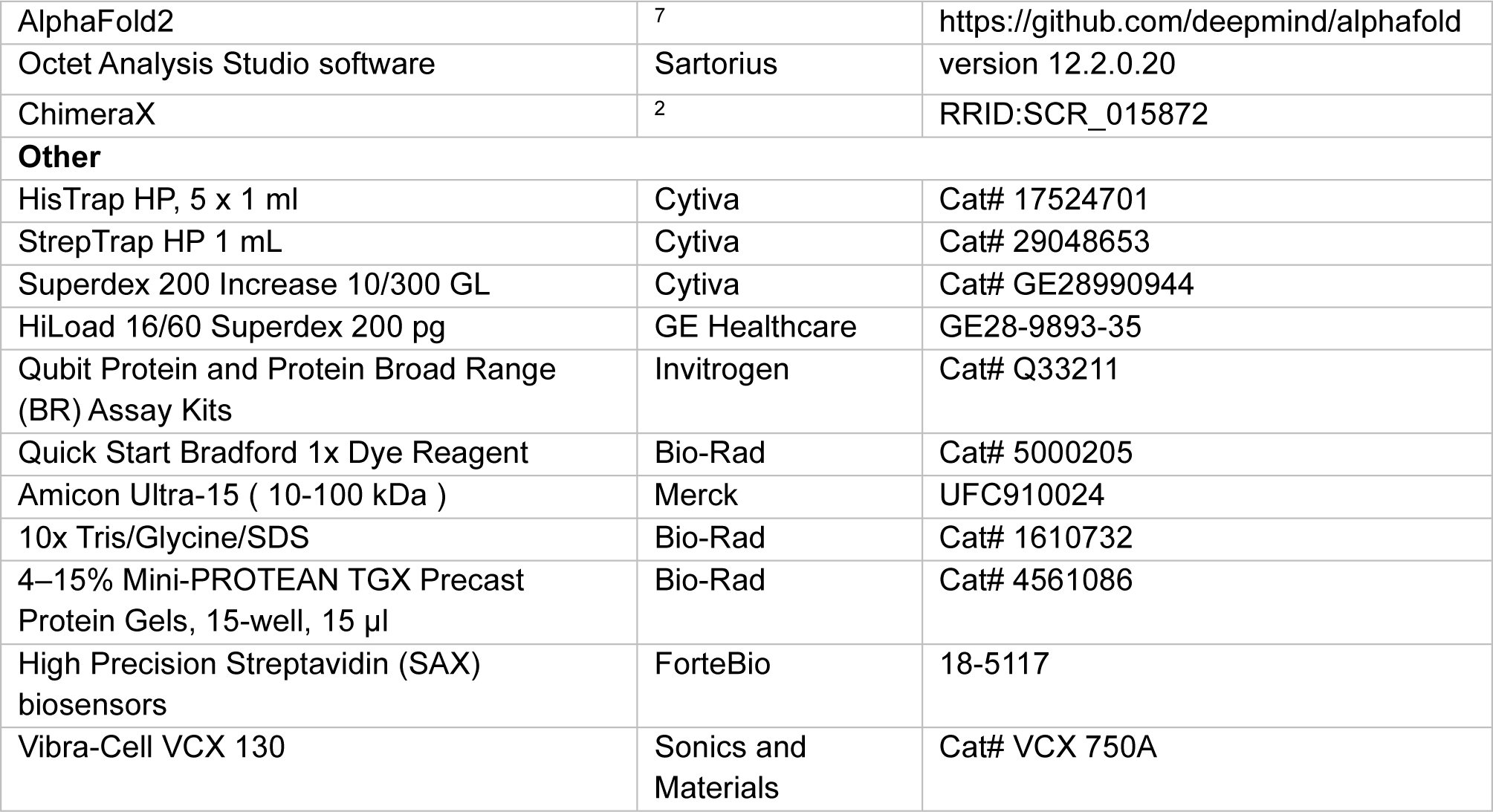
List of oligonucleotides and DNA substrates used in the study.

**Supplementary Table S3.** EM data collection and refinement statistics.

|  | **BrxX<sub>apo</sub>** | **BrxX:Ocr**<br>**(BrxX<sub>2</sub>:Ocr<sub>2</sub>)** | **BrxX:DNA**<br>**(43bp, H2T)** |
| --- | --- | --- | --- |
|  | PDB: 9EXH | PDB: 9EX7 | PDB: 9EWZ |
| **Data collection** |  |  |  |
| Microscope | TFS Krios | TFS Krios | TFS Krios |
| Magnification | 120 000 X | 120 000X | 165 000 X |
| Voltage (kV) | 300 | 300 | 300 |
| Electron dose (e<sup>-</sup>/ Å<sup>-2</sup>) | 35.1 | 35.1 | 50.5 |
| Detector | FEI Falcon IV (4k x 4k) | FEI Falcon IV (4k x 4k) | FEI Falcon IV (4k x 4k) |
| Defocus range (-μm) | 3 – 0.9 | 3 – 0.9 | 2.4 – 0.7 |
| Pixel size (Å) | 0.68 | 0.68 | 0.74 |
| Symmetry imposed | C1 | C1 | C1 |
| Micrographs (no.) | 5 100 | 5 100 | 11 152 |
| Initial particle images (no.) | 396 923 | 396 923 | 610 100 |
| Final particle images (no.) | 24 284 | 94 764 | 216 233 |
| Global map resolution (Å) | 4.13 | 2.84 | 2.22 |
| Fourier shell correlation (FSC) threshold | 0.143 | 0.143 | 0.143 |
| **Refinement Statistics** |  |  |  |
| Model Resolution (Å) | 4.0 (4.1) | 2.91 (3.44) | 2.21 (2.54) |
| FSC Threshold | 0.143 (0.5) | 0.143 (0.5) | 0.143 (0.5) |
| Map sharpening *B* factor (Å<sup>2</sup>) | -- | -- | -- |
| **Model Composition** |  |  |  |
| Non-hydrogen atoms | 9 728 | 21 099 | 10 789 |
| Protein residues | 1205 | 1205 | 1205 |
| Nucleotides | 0 | 0 | 50 |
| Ligands | 0 | 2 | 2 |
| ***B* factors (Å<sup>2</sup>)** |  |  |  |
| Protein | 183.81 | 132.03 | 93.98 |
| Nucleotides | 0 | 0 | 80.39 |
| Ligands | 0 | 96.36 | 76.84 |
| **R.M.S deviations** |  |  |  |
| Bond lengths (Å) | 0.002 | 0.003 | 0.002 |
| Bond angles (°) | 0.414 | 0.499 | 0.470 |
| **Validation** |  |  |  |
| MolProbity score | 1.67 | 1.98 | 1.40 |
| Clashscore | 7.78 | 4.78 | 3.54 |
| **Ramachandran statistics** |  |  |  |
| Favored (%) | 96.34 | 95.24 | 96.26 |
| Allowed (%) | 3.66 | 4.76 | 3.74 |
| Outliers (%) | 0.00 | 0.00 | 0.00 |

**Supplementary Table S4.** Structural alignments of MTD and TRD of BrxX (DNA-bound) with homologous enzymes, and with an Ocr-bound structure. Shown are outputs from ChimeraX *matchmaker* programme.

|  |  | CamA |  | MmeI |  | DrdV |  | HsdM |  | Ocr-Bound BrxX |  |  |  |
| --- | --- | --- | --- | --- | --- | --- | --- | --- | --- | --- | --- | --- | --- |
|  |  | MTD | TRD | MTD | TRD | MTD | TRD | MTD | TRD | NTD | MTD | TRD | CTD |
| **DNA-bound BrxX** | RMSD (Pruned Atom Pairs) | 1.056 (154) | 1.091 (72) | 1.113 (103) | 1.323 (17) | 0.988 (128) | 1.463 (6) | 1.101 (101) | 0.239 (9) | 0.891 (209) | 0.888 (392) | 0.838 (173) | 0.854 (339) |
|  | RMSD (All Pairs) | 13.295 (271) | 5.160 (164) | 11.816 (296) | 21.879 (159) | 8.358 (274) | 22.205 (70) | 12.980 (241) | 16.905 (40) | 0.901 (210) | 1.324 (416) | 1.085 (187) | 1.570 (368) |

## REFERENCES

1. Parikka, K. J., Le Romancer, M., Wauters, N. & Jacquet, S. Deciphering the virus-to-prokaryote ratio (VPR): insights into virus–host relationships in a variety of ecosystems. Biological reviews 92, 1081–1100 (2017).

2. Dion, M. B., Oechslin, F. & Moineau, S. Phage diversity, genomics and phylogeny. Nat Rev Microbiol 18>, 125–138 (2020).

3. Bernheim, A. & Sorek, R. The pan-immune system of bacteria: antiviral defence as a community resource. Nat Rev Microbiol 18, 113–119 (2020).

4. Mayo-Muñoz, D., Pinilla-Redondo, R., Birkholz, N. & Fineran, P. C. A host of armor: Prokaryotic immune strategies against mobile genetic elements. Cell Rep 42, (2023).

5. Miller, D., Stern, A. & Burstein, D. Deciphering microbial gene function using natural language processing. Nat Commun 13, 5731 (2022).

6. Koonin, E. V., Makarova, K. S. & Wolf, Y. I. Evolutionary Genomics of Defense Systems in Archaea and Bacteria. Annu Rev Microbiol 71, 233–261 (2017).

7. Isaev, A. B., Musharova, O. S. & Severinov, K. V. Microbial arsenal of antiviral defenses– part I. Biochemistry (Moscow) 86, 319–337 (2021).

8. Lopatina, A., Tal, N. & Sorek, R. Abortive infection: bacterial suicide as an antiviral immune strategy. Annu Rev Virol 7, 371–384 (2020).

9. LeRoux, M. & Laub, M. T. Toxin-antitoxin systems as phage defense elements. Annu Rev Microbiol 76, 21–43 (2022).

10. Tock, M. R. & Dryden, D. T. F. The biology of restriction and anti-restriction. 8, 466–472 (2005).

11. Tesson, F. et al. Systematic and quantitative view of the antiviral arsenal of prokaryotes. Nat Commun 13, 1–10 (2022).

12. Wang, L., Jiang, S., Deng, Z., Dedon, P. C. & Chen, S. DNA phosphorothioate modification—a new multi-functional epigenetic system in bacteria. FEMS Microbiol Rev 43, 109–122 (2019).

13. Xiong, X. et al. SspABCD–SspE is a phosphorothioation-sensing bacterial defence system with broad anti-phage activities. Nat Microbiol 5, 917–928 (2020).

14. Gedara, S. H. et al. 7-Deazaguanines in DNA: functional and structural elucidation of a DNA modification system. Nucleic Acids Res 51, 3836–3854 (2023).

15. Ofir, G. et al. DISARM is a widespread bacterial defence system with broad anti-phage activities. Nat Microbiol 3, 90–98 (2018).

16. Maestri, A. et al. Bacterial defences interact synergistically by disrupting phage cooperation. bioRxiv 2023 (2023).

17. Goldfarb, T. et al. BREX is a novel phage resistance system widespread in microbial genomes. EMBO J 34, 169–183 (2015).

18. Gordeeva, J. et al. BREX system of Escherichia coli distinguishes self from non-self by methylation of a specific DNA site. Nucleic Acids Res 47, 253–265 (2019).

19. Gao, H. et al. Nicking mechanism underlying the DNA phosphorothioate-sensing antiphage defense by SspE. Nat Commun 13, 6773 (2022).

20. Bravo, J. P. K. et al. Structural basis for broad anti-phage immunity by DISARM. Nat Commun 13, 1–11 (2022).

21. Makarova, K. S., Wolf, Y. I., Snir, S. & Koonin, E. V. Defense islands in bacterial and archaeal genomes and prediction of novel defense systems. J Bacteriol 193, 6039–56 (2011).

22. Chinenova, T. A., Mkrtumian, N. M. & Lomovskaia, N. D. Genetic characteristics of a new phage resistance trait in Streptomyces coelicolor A3 (2). Genetika 18, 1945–1952 (1982).

23. Hoskisson, P. A., Sumby, P. & Smith, M. C. M. The phage growth limitation system in Streptomyces coelicolor A(3)2 is a toxin/antitoxin system, comprising enzymes with DNA methyltransferase, protein kinase and ATPase activity. Virology 477, 100–109 (2015).

24. Rao, D. N., Dryden, D. T. F. & Bheemanaik, S. Type III restriction-modification enzymes: a historical perspective. Nucleic Acids Res 42, 45–55 (2014).

25. Morgan, R. D., Bhatia, T. K., Lovasco, L. & Davis, T. B. MmeI: a minimal Type II restriction-modification system that only modifies one DNA strand for host protection. Nucleic Acids Res 36, 6558–6570 (2008).

26. Kulkarni, M., Nirwan, N., van Aelst, K., Szczelkun, M. D. & Saikrishnan, K. Structural insights into DNA sequence recognition by Type ISP restriction-modification enzymes. Nucleic Acids Res 44, 4396–4408 (2016).

27. Gupta, Y. K., Chan, S.-H., Xu, S. & Aggarwal, A. K. Structural basis of asymmetric DNA methylation and ATP-triggered long-range diffusion by EcoP15I. Nat Commun 6, 1–10 (2015).

28. Chand, M. K. et al. Translocation-coupled DNA cleavage by the Type ISP restriction-modification enzymes. Nat Chem Biol 11, 870 (2015).

29. Shen, B. W. et al. Coordination of phage genome degradation versus host genome protection by a bifunctional restriction-modification enzyme visualized by CryoEM. Structure 29, 521– 530 (2021).

30. Hui, W. et al. Identification and functional analysis of the pglX gene of Lactobacillus casei Zhang bacteriophage exclusive (BREX) system. Appl Environ Microbiol AEM-01001 (2019).

31. Picton, D. M. et al. The phage defence island of a multidrug resistant plasmid uses both BREX and type IV restriction for complementary protection from viruses. Nucleic Acids Res (2021).

32. Zaworski, J. et al. Reassembling a cannon in the DNA defense arsenal: Genetics of StySA, a BREX phage exclusion system in Salmonella lab strains. PLoS Genet 18, e1009943 (2022).

33. Luyten, Y. A. et al. Identification and characterization of the WYL BrxR protein and its gene as separable regulatory elements of a BREX phage restriction system. Nucleic Acids Res 50, 5171–5190 (2022).

34. Wozniak, C. E., Hughes, K. T. & Liou, T. G. Mutations in the C-terminal region of the bacteriophage exclusion protein PglX can selectively inactivate restriction in Salmonella. J Bacteriol 205, e00207–23 (2023).

35. Schmitz, C. et al. Response regulator PorX coordinates oligonucleotide signalling and gene expression to control the secretion of virulence factors. Nucleic Acids Res 50, 12558–12577 (2022).

36. Greci, M. D. & Bell, S. D. Archaeal DNA replication. Annu Rev Microbiol 74, 65–80 (2020).

37. Attali, I., Botchan, M. R. & Berger, J. M. Structural mechanisms for replicating DNA in eukaryotes. Annu Rev Biochem 90, 77–106 (2021).

38. Shen, B. W. et al. Structure, substrate binding and activity of a unique AAA+ protein: the BrxL phage restriction factor. Nucleic Acids Res 51, 3513–3528 (2023).

39. Beck, I. N., Picton, D. M. & Blower, T. R. Crystal structure of the BREX phage defence protein BrxA. Curr Res Struct Biol 4, 211–219 (2022).

40. Picton, D. M. et al. A widespread family of WYL-domain transcriptional regulators co-localizes with diverse phage defence systems and islands. Nucleic Acids Res 50, 5191–5207 (2022).

41. Blankenchip, C. L. et al. Control of bacterial immune signaling by a WYL domain transcription factor. Nucleic Acids Res 50, 5239–5250 (2022).

42. Payne, L. J., Hughes, T. C. D., Fineran, P. C. & Jackson, S. A. New antiviral defences are genetically embedded within prokaryotic immune systems. bioRxiv 2021–2024 (2024).

43. Isaev, A. et al. Phage T7 DNA mimic protein Ocr is a potent inhibitor of BREX defence. Nucleic Acids Res 48, 5397–5406 (2020).

44. Atanasiu, C., Su, T. J., Sturrock, S. S. & Dryden, D. T. F. Interaction of the ocr gene 0.3 protein of bacteriophage T7 with EcoKl restriction/modification enzyme. Nucleic Acids Res 30, 3936–3944 (2002).

45. Sistla, S. & Rao, D. N. S-Adenosyl-L-methionine–dependent restriction enzymes. Crit Rev Biochem Mol Biol 39, 1–19 (2004).

46. Andriianov, A. et al. Phage T3 overcomes the BREX defense through SAM cleavage and inhibition of SAM synthesis by SAM lyase. Cell Rep 42, (2023).

47. Letellier, L., Plançon, L., Bonhivers, M. & Boulanger, P. Phage DNA transport across membranes. Res Microbiol 150, 499–505 (1999).

48. Cumby, N., Edwards, A. M., Davidson, A. R. & Maxwell, K. L. The bacteriophage HK97 gp15 moron element encodes a novel superinfection exclusion protein. J Bacteriol 194, 5012–5019 (2012).

49. Konttinen, O., Carmody, J., Kurnik, M., Johnson, K. A. & Reich, N. High fidelity DNA strand-separation is the major specificity determinant in DNA methyltransferase CcrM’s catalytic mechanism. Nucleic Acids Res gkad443 (2023).

50. Taylor, I., Watts, D. & Kneale, G. Substrate recognition and selectivity in the type IC DNA modification methylase M. EcoR124l. Nucleic Acids Res 21, 4929–4935 (1993).

51. Cheng, X. & Roberts, R. J. AdoMet-dependent methylation, DNA methyltransferases and base flipping. Nucleic Acids Res 29, 3784–3795 (2001).

52. Pettersen, E. F. et al. UCSF ChimeraX: Structure visualization for researchers, educators, and developers. Protein science 30, 70–82 (2021).

53. Malone, T., Blumenthal, R. M. & Cheng, X. Structure-guided analysis reveals nine sequence motifs conserved among DNA amino-methyl-transferases, and suggests a catalytic mechanism for these enzymes. J Mol Biol 253, 618–632 (1995).

54. Ashkenazy, H. et al. ConSurf 2016: an improved methodology to estimate and visualize evolutionary conservation in macromolecules. Nucleic Acids Res 44, W344–W350 (2016).

55. Holm, L. & Rosenstrï¿ ½m, P. ½ivi. Dali server: conservation mapping in 3D. Nucleic Acids Res 38, W545–W549 (2010).

56. Van Kempen, M. et al. Fast and accurate protein structure search with Foldseek. Nat Biotechnol 1–4 (2023).

57. Callahan, S. J. et al. Structure of type IIL restriction-modification enzyme MmeI in complex with DNA has implications for engineering new specificities. PLoS Biol 14, e1002442 (2016).

58. Zhou, J., Horton, J. R., Blumenthal, R. M., Zhang, X. & Cheng, X. Clostridioides difficile specific DNA adenine methyltransferase CamA squeezes and flips adenine out of DNA helix. Nat Commun 12, 3436 (2021).

59. Kennaway, C. K. et al. The structure of M. EcoKI Type I DNA methyltransferase with a DNA mimic antirestriction protein. Nucleic Acids Res 37, 762–770 (2009).

60. Quintana-Feliciano, R. et al. Burkholderia cenocepacia epigenetic regulator M. BceJIV simultaneously engages two DNA recognition sequences for methylation. bioRxiv 2021–2024 (2024).

61. Sam C. Went et al. Structure and rational engineering of the PglX methyltransferase and specificity factor for BREX phage defence. bioRxiv (2024).

62. Deep, A., Liang, Q., Enustun, E., Pogliano, J. & Corbett, K. D. Architecture and infection-sensing mechanism of the bacterial PARIS defense system. bioRxiv 2021–2024 (2024).

63. Burman, N. et al. Viral proteins activate PARIS-mediated tRNA degradation and viral tRNAs rescue infection. bioRxiv 2021–2024 (2024).

64. Bauerle, M. R., Schwalm, E. L. & Booker, S. J. Mechanistic diversity of radical S-adenosylmethionine (SAM)-dependent methylation. Journal of Biological Chemistry 290, 3995–4002 (2015).

65. Nelson, M., Raschke, E. & McClelland, M. Effect of site-specific methylation on restriction endonucleases and DNA modification methyltransferases. Nucleic Acids Res 21, 3139 (1993).

66. Samson, R. Y., Abeyrathne, P. D. & Bell, S. D. Mechanism of archaeal MCM helicase recruitment to DNA replication origins. Mol Cell 61, 287–296 (2016).

67. Frigola, J., Remus, D., Mehanna, A. & Diffley, J. F. X. ATPase-dependent quality control of DNA replication origin licensing. Nature 495, 339–343 (2013).

68. Srikhanta, Y. N., Fox, K. L. & Jennings, M. P. The phasevarion: phase variation of type III DNA methyltransferases controls coordinated switching in multiple genes. Nat Rev Microbiol 8, 196–206 (2010).

69. Punjani, A., Rubinstein, J. L., Fleet, D. J. & Brubaker, M. A. cryoSPARC: algorithms for rapid unsupervised cryo-EM structure determination. Nat Methods 14, 290–296 (2017).

70. Bepler, T. et al. Positive-unlabeled convolutional neural networks for particle picking in cryo-electron micrographs. Nat Methods 16, 1153–1160 (2019).

71. Punjani, A., Zhang, H. & Fleet, D. J. Non-uniform refinement: adaptive regularization improves single-particle cryo-EM reconstruction. Nat Methods 17, 1214–1221 (2020).

72. Zivanov, J., Nakane, T. & Scheres, S. H. W. A Bayesian approach to beam-induced motion correction in cryo-EM single-particle analysis. IUCrJ 6, 5–17 (2019).

73. Zivanov, J., Nakane, T. & Scheres, S. H. W. Estimation of high-order aberrations and anisotropic magnification from cryo-EM data sets in RELION-3.1. IUCrJ 7, 253–267 (2020).

74. Wolf, M., DeRosier, D. J. & Grigorieff, N. Ewald sphere correction for single-particle electron microscopy. Ultramicroscopy 106, 376–382 (2006).

75. Emsley, P., Lohkamp, B., Scott, W. G. & Cowtan, K. Features and development of Coot. Acta Crystallogr D Biol Crystallogr 66, 486–501 (2010).

76. Wang, X., Terashi, G. & Kihara, D. CryoREAD: de novo structure modeling for nucleic acids in cryo-EM maps using deep learning. Nat Methods 20, 1739–1747 (2023).

77. Liebschner, D. et al. Macromolecular structure determination using X-rays, neutrons and electrons: recent developments in Phenix. Acta Crystallogr D Struct Biol 75, 861–877 (2019).

78. Williams, C. J. et al. MolProbity: More and better reference data for improved all-atom structure validation. Protein Science 27, 293–315 (2018).

79. Afonine, P. V et al. New tools for the analysis and validation of cryo-EM maps and atomic models. Acta Crystallogr D Struct Biol 74, 814–840 (2018).

80. Sambrook, J. & Russell, D. W. Purification of nucleic acids by extraction with phenol: chloroform. Cold Spring Harb Protoc 2006, pdb-prot4455 (2006).

81. Clokie, M. R. J., Kropinski, A. M. & Lavigne, R. Bacteriophages. vol. 501 (Springer, 2009).

## SUPPLEMENTARY REFERENCES

1. Ashkenazy, H., Abadi, S., Martz, E., Chay, O., Mayrose, I., Pupko, T., and Ben-Tal, N. (2016). ConSurf 2016: an improved methodology to estimate and visualize evolutionary conservation in macromolecules. Nucleic Acids Res 44, W344–W350.

2. Pettersen, E.F., Goddard, T.D., Huang, C.C., Meng, E.C., Couch, G.S., Croll, T.I., Morris, J.H., and Ferrin, T.E. (2021). UCSF ChimeraX: Structure visualization for researchers, educators, and developers. Protein science 30, 70–82.

3. Gordeeva, J., Morozova, N., Sierro, N., Isaev, A., Sinkunas, T., Tsvetkova, K., Matlashov, M., Truncaitė, L., Morgan, R.D., and Ivanov, N. V (2019). BREX system of Escherichia coli distinguishes self from non-self by methylation of a specific DNA site. Nucleic Acids Res 47, 253–265.

4. Isaev, A., Drobiazko, A., Sierro, N., Gordeeva, J., Yosef, I., Qimron, U., Ivanov, N. V, and Severinov, K. (2020). Phage T7 DNA mimic protein Ocr is a potent inhibitor of BREX defence. Nucleic Acids Res 48, 5397–5406.

5. Liebschner, D., Afonine, P. V, Baker, M.L., Bunkóczi, G., Chen, V.B., Croll, T.I., Hintze, B., Hung, L.-W., Jain, S., and McCoy, A.J. (2019). Macromolecular structure determination using X-rays, neutrons and electrons: recent developments in Phenix. Acta Crystallogr D Struct Biol 75, 861–877.

6. Punjani, A., Rubinstein, J.L., Fleet, D.J., and Brubaker, M.A. (2017). cryoSPARC: algorithms for rapid unsupervised cryo-EM structure determination. Nat Methods 14, 290–296.

7. Jumper, J., Evans, R., Pritzel, A., Green, T., Figurnov, M., Ronneberger, O., Tunyasuvunakool, K., Bates, R., Žídek, A., and Potapenko, A. (2021). Highly accurate protein structure prediction with AlphaFold. Nature 596, 583–589.

